# Electroadhesive hydrogel interface for prolonged mucosal theranostics

**DOI:** 10.1101/2023.12.19.572448

**Authors:** Binbin Ying, Kewang Nan, Qing Zhu, Tom Khuu, Hana Ro, Sophia Qin, Shubing Wang, Karen Jiang, Yonglin Chen, Guangyu Bao, Josh Jenkins, Andrew Pettinari, Johannes Kuosmanen, Keiko Ishida, Niora Fabian, Aaron Lopes, Jason Li, Alison Hayward, Robert Langer, Giovanni Traverso

## Abstract

Establishing a robust and intimate mucosal interface that allows medical devices to remain within lumen-confined organs for extended periods has valuable applications, particularly for gastrointestinal (GI) theranostics. Here, we report the development of **e-GLUE**, an **e**lectroadhesive hydro**g**e**l** interface for robust and prolonged m**u**cosal r**e**tention following electrical activation. Notably, this novel mucosal adhesion mechanism can increase the adhesion energy of hydrogels on the mucosa by up to 30-fold and enable *in vivo* GI retention of e-GLUE devices for up to 30 days. Strong mucosal adhesion occurs within one minute of electrical activation, despite the presence of luminal fluid, mucus exposure, and organ motility, thereby ensuring compatibility with complex in vivo environments. In swine studies, we demonstrate the utility of e-GLUE for mucosal hemostasis, sustained local delivery of therapeutics, and intimate biosensing in the GI tract. This system can enable improved treatments for various health conditions, including gastrointestinal bleeding, inflammatory bowel disease, and diagnostic applications in the GI tract and beyond.

## Main Text

### Introduction

Mucosa adhesives are valuable for sustained biomedical applications such as hemostatic sealing (*1*), drug delivery (*2*), and physiological sensing (*3*), targeting specific regions in lumen-confined organs such as gastrointestinal (GI) tract (**Fig. 1A**). However, existing adhesion mechanisms can generally be retained in the GI tract for up to 24 hours owing to the complex GI biochemical environment, rapid turnover of mucus and surface epithelium, and constant gastrointestinal motility (*4*–*6*). For example, the lifetime of currently available mucoadhesives (*7*) and hydrogel tissue adhesives (*8, 9*) is approximately 24 hours or less in the GI tract, as physical and chemical bonding sites solely form in the mucus layers (**fig. S1A, B**), which are rapidly eliminated through mucociliary clearance within hours (*10*). Their adhesion ability is further attenuated by gastric fluid (*9*). To enhance mucosal retention time, adhesives should be able to penetrate the mucosal layers and interact with the deep mucosal epithelium comprised of cell types with a much slower turnover rate (*11*). Topological adhesion that enables physical entanglement with tissues through polymer penetration (*12, 13*) can only reach the outermost mucosal layer after 24 hours due to the slow speed of passive diffusion (**fig. S1C**). While clinical tools (e.g., endoclips) can rapidly hook onto the GI mucosa for weeks to months, these approaches are usually traumatic (*14, 15*). Here, we developed a novel adhesion mechanism that can interact with deep mucosal epithelium to prolong mucosal retention for up to 30 days in the GI tract. This new mechanism could enable novel applications including treatments for recurrent GI bleeding (*16*), long-term and localized therapeutics (*17*), and biosensing devices requiring intimate sensor-mucosa interaction (*18, 19*).

**Fig. 1.**
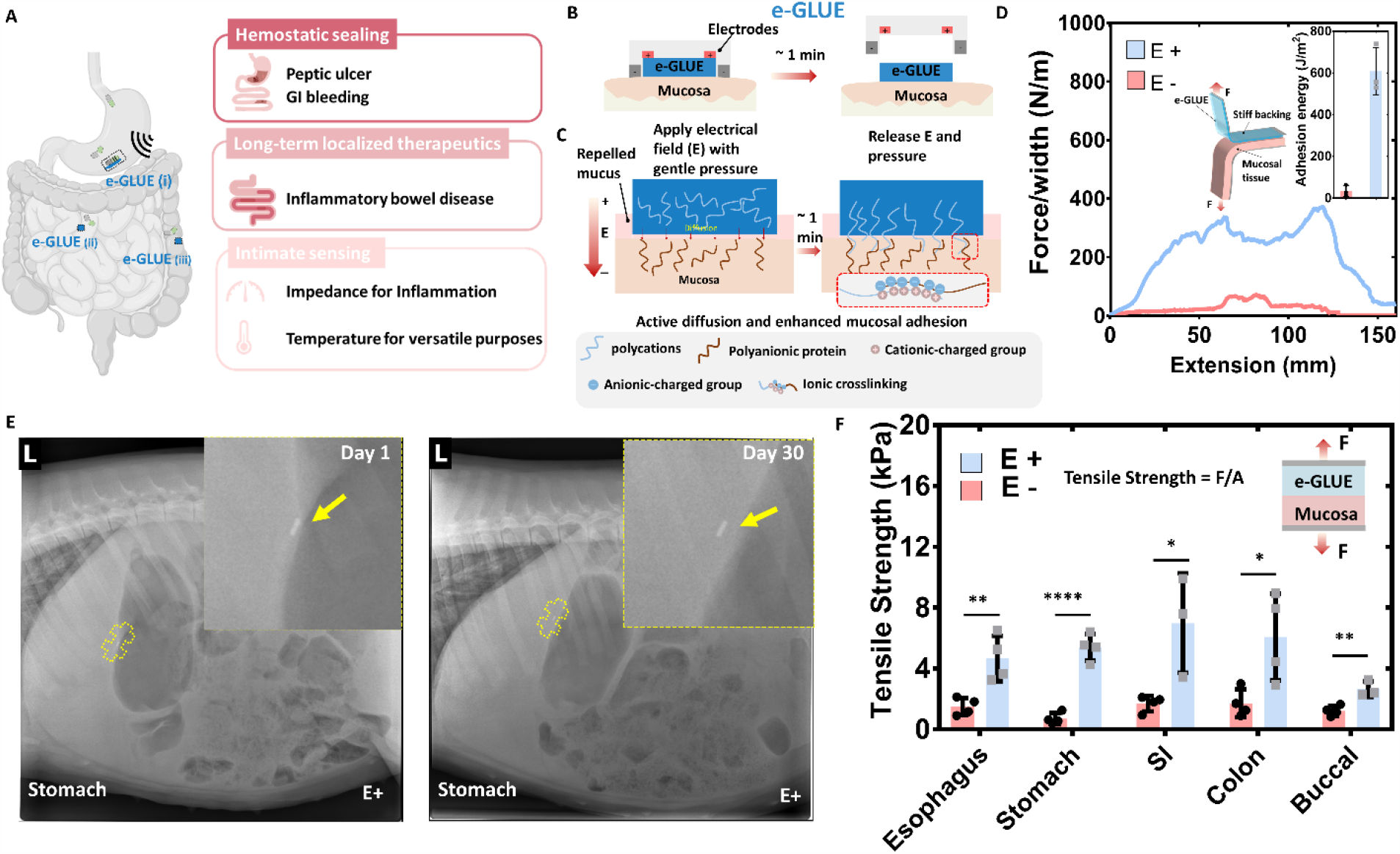
Electroadhesion interaction between mucosal tissues and e-GLUE. **A)** Diverse gastrointestinal conditions and theranostic opportunities. **B)** Schematic illustration of e-GLUE deployment with electrodes on the mucosal surface. **C**) The electroadhesive mechanism responsible for the strong bonding between e-GLUE and mucosal tissues relies on a combination of active polymeric diffusion and polycationic-polyanionic crosslinking. **D**) Representative force-extension curves of e-GULE mucosa hybrids with electrical stimulation treatment (**E+**) or without (**E-**) during peeling tests. Inset indicates their adhesion energy difference. **E**) X-ray images show that e-GLUE retention time on the gastric mucosa can be over 30 days (n=3). Yellow arrows show the gastric retention location of e-GLUE with X-ray opaque beads. **F**) Tensile strength of various GI mucosal tissues adhered by e-GLUE with **(E+**) and without (**E-**) electrical stimulation (15 V, 40 s). Square-shaped electrodes were used for non-invasive bonding. e-GLUE thickness is 0.75 mm, and polycations concentration is 10%. SI, small intestinal. The values in panels **D** and **F** indicate the mean and the standard deviation (n = 3–4). Statistical significance and corresponding P values were determined by a two-sided student’s t-test. *P < 0.05; **P < 0.01; ****P < 0.0001.

We report the development of **e-GLUE**, an **e**lectroadhesive hydro**g**e**l** interface for robust and prolonged m**u**cosal r**e**tention (**Fig. 1A**). e-GLUE consists of cationic polymers interpenetrated in a tough hydrogel matrix. With an electrical stimulation treatment (∼1 min), cationic polymers can interact with polyanionic proteins within the mucosal tissue (**Fig. 1B** and **C)** that have a relatively slow cellular turnover rate (e.g., >7 days in GI tract) (*11*). This novel mechanism increases the adhesion energy between the hydrogel and mucosa by up to 30-fold (*E+*, **Fig. 1D, Movie S1**) and extends *in vivo* GI retention of e-GLUE devices for over 30 days through intimate mucosal adhesion (**Fig. 1E**). The rapid and robust adhesion between e-GLUE and the mucosa under electrical stimulation leverages a combination of active polymeric diffusion and polycationic-polyanionic crosslinking (**Fig. 1C**). To validate the adhesion efficacy, we performed electroadhesion throughout the entire mucosal tissues including the GI tract and observed a significant increase in adhesion strength (**Fig. 1F**), despite the presence of intraluminal fluid and mucus layers. We demonstrated several potential applications for e-GLUE, including instant mucosal homeostasis, prolonged local delivery of therapeutics, and intimate biosensing in the GI tract, through *in vivo* swine studies (**Fig. 1A**).

## Results

### e-GLUE design and electroadhesion mechanism

We designed e-GLUE as a tough hydrogel composed of polycations. Tough hydrogels have been widely utilized for various biomedical applications due to their tissue-like strength, stretchability, and biocompatibility (*20*). We selected polycations due to their electrostatic interaction with polyanionic proteins in the mucosal tissues (*21*). A tough hydrogel matrix was chosen to be compatible with polycations; both positive-charged double-network hydrogels [e.g., chitosan-polyacrylamide (Chi-PAAm) (*22*)] and neutral ones [e.g., agarose-PAAm hydrogels (*23*)] were considered. Unconventional neutral single-network hydrogels, such as highly entangled PAAm (*24*) or poly(ethylene glycol) (PEG) (*25*), were also considered. To identify appropriate polycations, we tested six types of polycations: PDAC, P3ATAC, P3METAC, P2ATAC, P2DAMA, and PN3DMA (**fig. S2**) with doping in a PAAm single-network hydrogel system. We chose alginate-PAAm (Alg-PAAm) hydrogel as a model mucosal tissue, as it closely resembles mucosa in terms of mechanical toughness and polyanionic properties (*20*). After applying a DC electrical stimulation on the e-GLUE mucosa hybrids (**fig. S3A**), we quantified their adhesion performance using 180° peel tests (**Fig. 1D**) and tensile tests (**fig. S3B**). Among the tested polycations, PDAC, P3ATAC, P3METAC, and P2ATAC were measured to have relatively higher adhesion strengths at >50 kPa (**Fig. 2A**), likely due to quaternary amine groups present on these polymers (**fig. S2**). For subsequent experiments, we doped PDAC, a commercially available polycation, into the Chi-PAAm hydrogel, resulting in a PDAC-based e-GLUE (**fig. S4A, B**) (see **Methods** for the synthesis detail). The resulting e-GLUE exhibited a *Young’s* modulus of 13.6 kPa (**fig. S4C**), which is 50∼100 times softer than the digestive tract (*26*). The flexibility of this PDAC-based e-GLUE could potentially relieve the mechanical constraint on regular organ motility (e.g., GI peristalsis) once the e-GLUE adheres to mucosal surfaces. To assess the biocompatibility of e-GLUE, we conducted *in vitro* cytotoxicity tests on cultured mouse embryonic fibroblast cells (NIH/3T3). LIVE/DEAD staining and MTT viability assays indicated that 78%± 2% (n=7) of the cells remained alive after 24 hours of exposure to the e-GLUE extraction (**fig. S5**), confirming its low cytotoxicity.

**Fig. 2.**
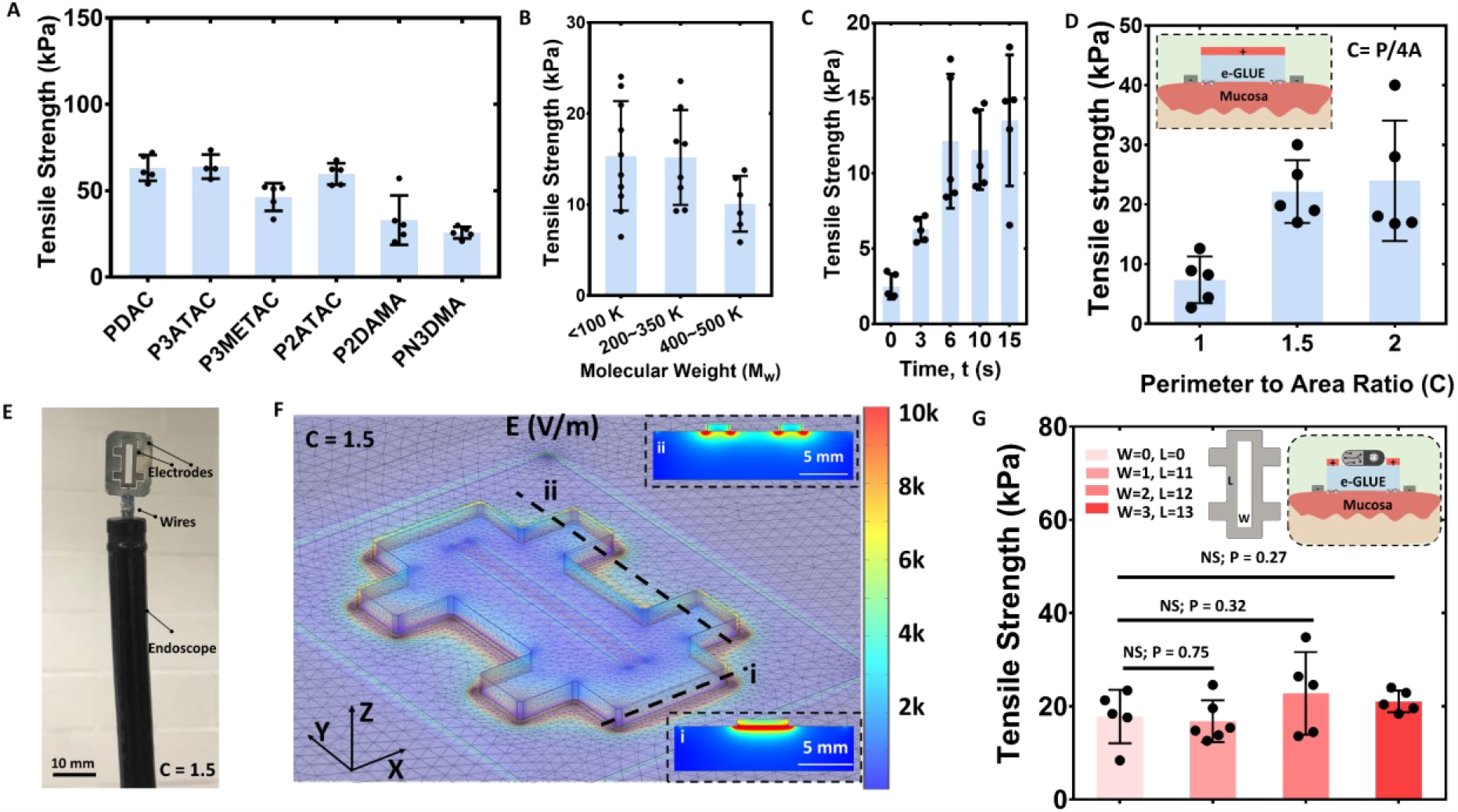
e-GLUE’s adhesion performance. The adhesion strength of e-GLUE depends on **A**) the type of polycation, **B**) the molecular weight or length of the cationic polymer chains (Mw), **C**) the electrical stimulation time (t), and three other factors shown in the supplementary material. **D**) Optimization of e-GLUE electrodes enables non-invasive deployment of e-GLUE on the mucosa. The tensile strength between jejunum mucosa and e-GLUE can be adjusted by varying the electrode’s perimeter (P) to area (A) ratio (C), where C = P/4A. The inset schematic provides the cross-sectional view of non-invasive e-GLUE electrodes. Values in panels **A-D** indicate the mean and the standard deviation (n = 5–10). **E**) A photo of e-GLUE electrodes (C= 1.5) inserted into the endoscope for e-GLUE deployment in the GI tract. The rectangle cutout size is set as 2*12 mm^2^, as detailed in **Fig. 2G. F)** Numerical modeling of the electrical field distribution of the novel e-GLUE electrode design (C= 1.5). The voltage amplitude was set at 10 V. Insets to visualize the electrical field distribution of two cross-sections. **G**). Adhesion strength between e-GLUE and duodenal mucosa under different rectangle cutout sizes. The inset schematic proposes a design for a cutout structure in the positive electrode, which can be used to bridge ingestible devices with mucosal tissues through e-GLUE. Values in panel **G** indicate the mean and the standard deviation (n = 5-6). Statistical significance and corresponding P-values were determined by a two-sided student’s t-test. Condition used for (**A**): d= 0.75 mm, U=15 V, t=40 s, ρ=5%, and Mw=200∼350 K; for (**B**): d= 0.75 mm, U=6 V, t=10 s, ρ=10 %; for (**C**): d= 0.75 mm, U=6 V, ρ=10 %, and Mw=200∼350 K; for (**D**) d= 1 mm, U=15 V, ρ=10 %, t= 40 s, Mw=200∼350 K; for (**G**) d= 0.75 mm, U=15 V, ρ=10 %, t= 40 s, Mw=200∼350 K.

We next investigated the adhesion mechanism between the e-GLUE and mucosa. We hypothesized that the strong bonding between e-GLUE and mucosal tissues that occurs after a short electrical stimulation results from a combination of active polymeric diffusion and polycationic-polyanionic crosslinking (*27*). To visualize the active polymeric diffusion effect, we conjugated fluorescein (FITC) on the polycationic chain in e-GLUE. We observed that the polycations barely penetrate mucosa without electrical stimulation (E-, **fig. S6A**). Longer stimulation time led to a greater polycationic diffusion depth until equilibrium after 60 s (∼80 μm, E+, **fig. S6A** and **B**). To validate polycationic-polyanionic crosslinking, we applied chitosan-free e-GLUE on the mucosal model and PAAm hydrogel, respectively. Electrical stimulation resulted in a 3-fold increase in the adhesion force between e-GLUE and the tissue model compared to the e-GLUE and PAAm hydrogel, indicating the importance of polyanions in the mucosal layer for crosslinking (**fig. S6C**). Furthermore, the physical entanglement effect between e-GLUE and mucosal layers was negligible, as no significant adhesion enhancement was observed between e-GLUE and PAAm before or after electrical stimulation (**fig. S6D**).

We then optimized electroadhesion conditions on the model mucosal tissue by modifying five parameters: e-GLUE thickness (*d*), polycation concentration (*ρ*), polycationic chain length *(Mw)*, voltage amplitude (*U*), and electrical stimulation time (*t*). Among the tested conditions, we observed that adhesion performance was negatively correlated with gel thickness (**fig. S7A**) and polycationic chain length (**Fig. 2B**), but positively correlated with voltage amplitude (**fig. S7B**), stimulation time (**Fig. 2C**), and polycation concentration (**fig. S7B**). Reducing thickness below 0.75 mm, increasing voltage amplitude above 6 V, and extending stimulation time beyond 6 s did not result in a significant increase in adhesion performance (**Fig. 2C, fig. S7A**, and **B**). These observations can be attributed to the saturation of the electrophoresis effect (*28*). Thus, we selected an e-GLUE with a thickness of 0.75 mm and a PDAC concentration of 10% for subsequent *ex vivo* and *in vivo* tests to study how the electrode design and stimulation conditions would affect mucosal electroadhesion and safety.

### e-GLUE electrode optimization

To support endoscopic-based translation of e-GLUE for GI tract application and beyond, we designed a novel endoscopically compatible e-GLUE electrode assembly that enables electrodes to be placed directly on the mucosal surface (**Fig. 1B**, inset in **Fig. 2D**, and **Fig. 2E**). This design eliminates the requirement for invasive submucosal placement of ground electrodes for electrical stimulation (**fig. S3A**). We manufactured square-shaped electrodes (**fig. S7D**) through laser cutting and arranged the electrode assemblies to enable non-invasive electrical stimulation. After applying electrical stimulation, we measured a significant increase in adhesion strength between the e-GLUE and the mucosal tissues throughout the entire GI tract (*E+*, **Fig. 1F**) as well as other mucosal surfaces in the urinary, female reproductive, and respiratory tracts (*E+*, **fig. S8)**, despite the presence of luminal fluid and mucus layers. In contrast, when electrical stimulation was not applied, the adhesion strength between the e-GLUE and tissue was significantly lower (*E-*, **Fig. 1F** and **fig. S8**). These results suggest that the e-GLUE technology can significantly increase the adhesion strength of hydrogels to various types of mucosal tissue.

We optimized the e-GLUE mucosal adhesion strength by adjusting the dimensions of the electrodes, specifically the perimeter (*P*) to area (*A*) ratio (*C=P/4A*). By increasing *C* from 1 to 2 (**Fig. 2D** and **fig. S7D**), the adhesion strength between jejunum mucosa and e-GLUE can be increased from 7.4 to 24 kPa (**Fig. 2D**). We developed a numerical electromagnetic simulation using COMSOL Multiphysics (version 6.0, **Extended text)** to visualize the electrical field distribution with different electrode shapes (**Fig. 2F, fig. S9**). Numerical results showed that our novel electrode configuration induced a concentrated electrical field surrounding the area between the positive (located on the e-GLUE) and ground (located on the mucosa) electrodes (**fig. S9)**. A larger *C* (e.g., 1.5, **Fig. 2E**) yielded a longer concentrated electrical field perimeter (**Fig. 2F**), resulting in a higher adhesion strength (e.g., 22.2 kPa, **Fig. 2D**). Increasing *C* beyond 1.5 did not significantly improve adhesion strength but involved electrode shape complexity (*C= 2*, **Fig. 2D, fig. S7D)**. In addition, our numerical model predicted there would be a low electrical field distribution in the central section of the e-GLUE (**fig. S9)**, which is consistent with the low adhesion strength observed during tissue tests (**Movie S2**). We also used the numerical model to optimize electrode design. By reducing the *gap* between the ground and positive electrodes from 3 mm to 0.5 mm, the width of the concentrated electrical field expanded from 0.094 mm (**fig. S10-v)** to 0.451 mm (**fig. S10-iii)**. A larger concentrated electrical field area would lead to a higher adhesion force between the e-GLUE and tissue, which was observed during *ex vivo* tests. Further *gap* reduction might not effectively improve the adhesion performance because the numerical model predicted no further increase in the width of the concentrated electrical field. (**fig. S10-ii, iii)**.

To enable ingestible devices to interface with mucosal tissues, we incorporated cutout structures in the positive electrode so that devices can adhere to the top/bottom of the e-GLUE for prolonged and intimate theranostic applications (inset in **Fig. 2G**). To study how the cutout geometry affects the adhesion performance, we compared rectangle cutouts with varied widths (*W*) and lengths (*L*). Among the tested conditions, the tensile strength between e-GLUE and duodenal mucosa shows no significant difference (**Fig. 2G**). Thus, an optimized e-GLUE electrode (*C =1*.*5, gap* = 0.5 mm) with cutout structures in the positive electrode (e.g., *2 mm * 12 mm*, **Fig. 2E, fig. S7E**) was selected for subsequent *in vivo* tests to study how electrical stimulation conditions would affect mucosal electroadhesion and safety.

### *In vivo* assessment of e-GLUE’s safety and gastric retention time

We then assessed the safety and long-term retention efficacy of e-GLUE on the GI mucosa using an *in vivo* model in swine. Electrical stimulation treatment enables strong bonding between e-GLUE and mucosal tissues but results in increased tissue temperature due to Joule heating. To identify a safe electrical stimulation range, we first used the optimized e-GLUE electrode (**Fig. 2E**) to investigate how two primary factors - voltage amplitude (*U*) and time (*t*) - affect tissue safety *in vivo* without sacrificing adhesion performance. On-site tensile tests were performed to quantify the adhesion strength between e-GLUE and mucosal tissues (e.g., SI and stomach). During the stimulation, we used an IR camera to monitor the mucosal tissue temperature of an anesthetized swine in the terminal experiments (**Fig. 3B**). Our tests showed that *U=10 V* and *t=80 s* yielded an optimal adhesive strength of 16∼17 kPa with the maximum tissue temperature of 43 °C (**Fig. 3A-C**). Increasing the stimulation time may not result in a higher adhesive strength (**fig. S11**) due to the electrophoresis saturation effect. Increasing voltage can reduce the time to achieve a comparable adhesion strength (20 V, 25 s in **Fig. 3A**) but induces a higher temperature rise (20 V, 25 s in **Fig. 3B**) as a result of a larger corresponding current and Joule heating (**fig. S12A)**.

**Fig. 3.**
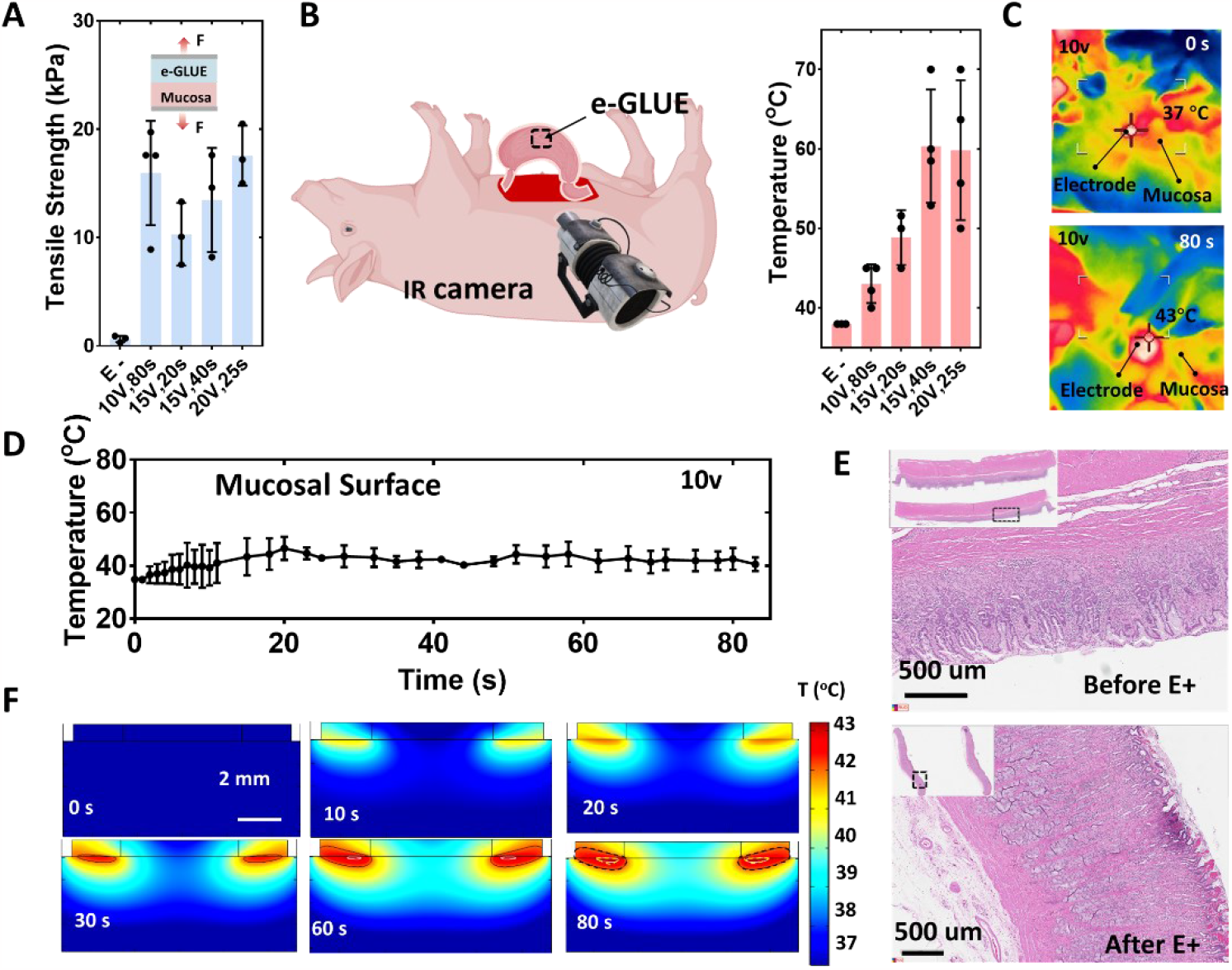
Optimization of electrical stimulation conditions and safety assessment for e-GLUE. **A**) e-GLUE’s adhesion strength on gastric mucosal tissue in vivo under various electrical stimulation conditions. **B**) Maximum temperature of local gastric tissue under various electrical stimulation conditions. The temperature was measured using an IR camera during terminal experiments where the pig was under anesthesia. **C**) IR images illustrating temperature distribution on the mucosal tissue before (0 s) and after (80 s) electrical stimulation treatment at a voltage amplitude of 10 V. **D**) Maximum temperature of local mucosal tissue over 80 s. **E**) Representative histological images of gastric mucosal tissues before and after electrical stimulation (E+) under optimal conditions (10 V, 80 s). Dashed rectangles indicate the regions around the e-GLUE’s perimeter. **F**) 3D electromagnetic-bioheat transfer simulations using COMSOL Multiphysics (version 6.0) to visualize spatial-temporal temperature distribution in the hydrogel-tissue coupling structure in the depth direction over 80 s. The voltage amplitude in **Fig. C, D**, and **F** was set at 10 V. Values in panels **A, B**, and **D** represent the mean and the standard deviation (n = 3-4).

Histological assessment by a pathologist (blinded to the status of the samples) indicated that at temperatures under 43 °C for 1 min (**Fig. 3D)**, the gastric mucosa exhibited mild inflammation around the e-GLUE’s perimeter (**Fig. 3E)** with no visible tissue damage (**fig S12B)**. Additionally, we included a bioheat module into the numerical simulations model (**Extended text**) to visualize the spatial-temporal temperature distribution of the coupled hydrogel-mucosa structure in the depth direction (**Fig. 3F)**. The simulation predicted that, with electrical stimulation, the surface mucosal tissue along the e-GLUE’s perimeter would heat to 43 °C by 20 s; due to the heat transfer effect, the top 200 μm of mucosal tissue would reach 43 °C by 80 s (red zones in **Fig. 3F, Movie S3)**. This numerical simulation is consistent with the histological assessment and temperature readings (**Fig. 3D** and **E**), supporting the safety of e-GLUE technology (*29*). In fact, the temperature increase can be further mitigated by applying pulse waveforms (**fig. S13A**) to dissipate heat without sacrificing adhesion strength (**fig. S13B**). Therefore, subsequent adhesion experiments using *U=10 V* and *t=80 s* for *in vivo* tests unless stated elsewhere.

To assess the long-term retention of e-GLUE on the GI mucosa, we conducted an endoscopic deployment by inserting the electrodes and e-GLUE through the esophagus into the swine’s stomach **(Fig. 4A 1-ii, fig. S14**, and **Movie S4)**. The GI tract exhibits constant motility, accompanied by rapid mucus turnover (∼12 hours) and mucosal epithelium cellular renewal (∼4 days) (*30*). As a result, achieving long-term retention within the dynamic and active GI luminal environment poses a considerable challenge. To visualize the gastric retention time, we embedded X-ray opaque stainless-steel rods into the e-GLUE. During the *in vivo* experiments, the gentle pressure from the endoscope ensured firm contact between the devices and the tissue in the dynamic GI environment (**Movie S3)**. After electrical stimulation, the electrodes, along with the endoscope, were retrieved from the GI tract (**Fig. 4A iii, fig. S14**). X-ray imaging and endoscopic examination revealed that e-GLUE’s retention time on the gastric mucosa ranges from 11 to over 30 days **(Fig. 1E** and **Fig. 4B)**, and it remained over 4 days on the colon mucosa **(Fig. 4B)**. In contrast, hydrogel tissue adhesives that mainly rely on physical and/or chemical bonding such as *N*-Hydroxysuccinimide (NHS) ester (*31*) were observed to pass through the GI tract within two days **(Fig. 4B** and **fig. S15)**. These results demonstrate the longer gastric retention time of e-GLUE compared to the existing adhesive strategies used on the GI tract (*32*–*34*). To assess the long-term safety of e-GLUE technology, mucosal tissues, including the duodenum, colon, and stomach, were examined 7-60 days after e-GLUE treatment; no scarring or inflammation was observed **(fig. S16A** and **B)**.

**Fig. 4.**
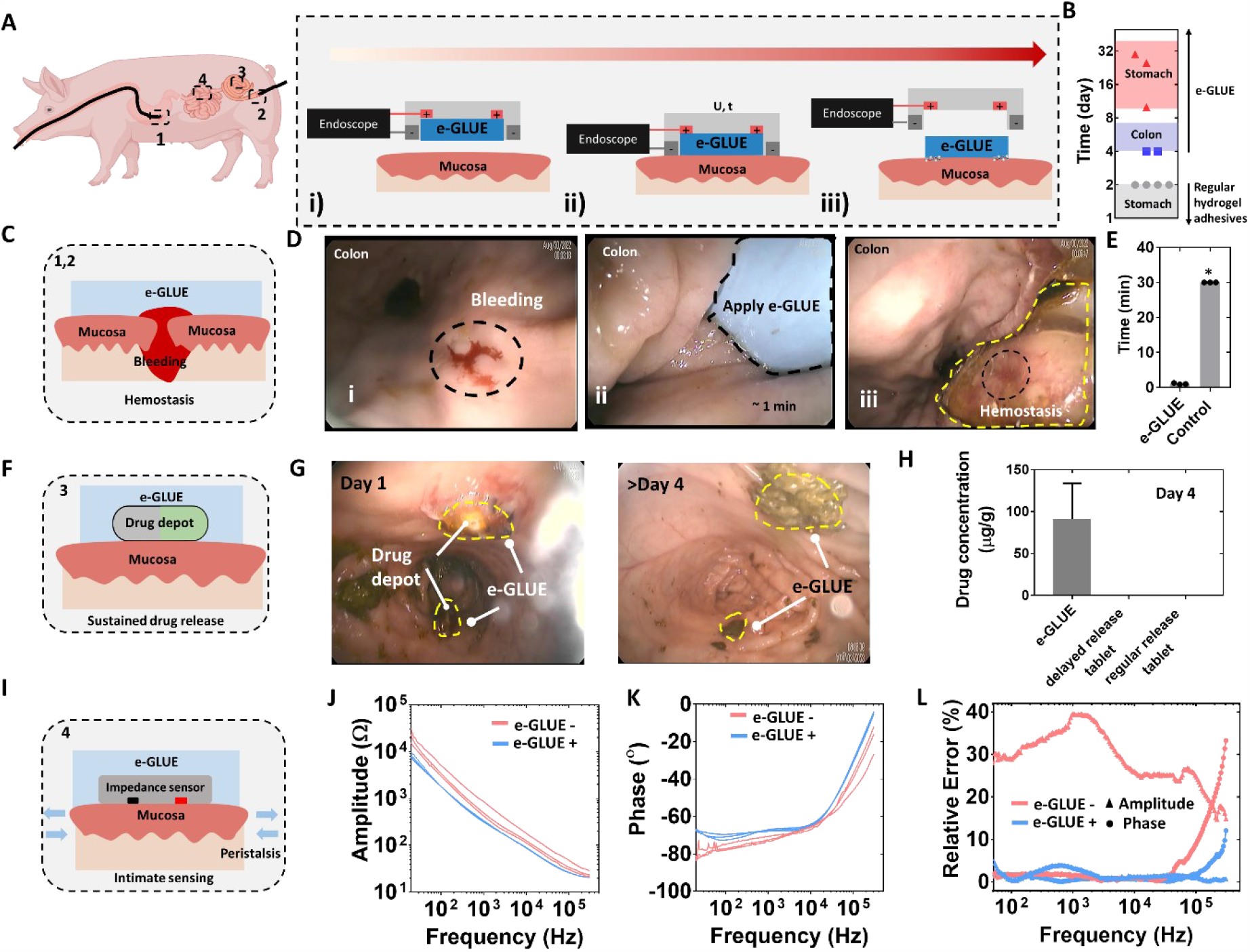
In vivo theranostic demonstration of e-GLUE. **A**) Schematic illustrating the endoscopic deployment of e-GLUE throughout the entire GI tract. **B)** Retention range of e-GLUE (red for gastric, blue for colon mucosa) and regular hydrogel adhesives (gray for gastric mucosa). Shapes represent the last visualization date under X-ray or endoscope. **C)** Schematic demonstrating e-GLUE’s capability for mucosal hemostasis. **D)** Endoscopic view visualizing the hemostasis procedures in the colon. **E**). Comparison of wound sealing time with and without e-GLUE on the gastric mucosa. *Untreated wounds on the stomach bled continuously for the entire 30-minute monitoring period. **F)** Budesonide-loaded e-GLUE adhered to mucosal tissues for prolonged local drug delivery. **G)** Endoscopic view showing the budesonide -loaded e-GLUE adhered to the colon mucosa for more than 4 days. **H)** Budesonide concentration on Day 4 in colon tissue through either e-GLUE or oral administration. **I**) Schematic illustrating the adhesion of an e-GLUE-sensor hybrid on mucosal tissues under peristalsis. **J**) Amplitude values and **K**) Phase values of three sets of impedance measurements on the same SI mucosal tissue using commercial sensors. **L**) Calculation of relative errors as the standard deviation divided by the mean value for sensors with e-GLUE (e-GLUE+) and without e-GLUE (e-GLUE-). Values in panel **E** and **H** represent the mean and the standard deviation (n = 3-4).

### Theranostic applications of e-GLUE

With both increased adhesion strength and extended gastric retention validated in vivo, we demonstrated e-GLUE for instant mucosal sealing, sustained local delivery of therapeutics, and intimate biosensing applications in the GI tract. GI bleeding and perforations are clinically life-threatening conditions (*35*). Here we report the e-GLUE system which can be used for instant hemostasis and sealing on the GI mucosa (**Fig. 4C**). To test this capability, we induced acute bleeding on the GI mucosa [e.g., colon (**Fig. 4D-i**) and stomach (**fig. S17A**)] in an anesthetized swine. Immediate e-GLUE treatment was applied on mucosal bleeding sites for ∼1 min (**Fig. 4D-ii**) with effective hemostasis observed (**Fig. 4D-iii, Fig. 4E, fig. S17B** and **C)**. In contrast, untreated wounds were observed to bleed continuously for the entire 30-minute monitoring period (**Fig. 4E**). With demonstrated ultra-prolonged adhesion time (**Fig. 1E** and **Fig. 4B**), our e-GLUE has the potential to eliminate recurrent GI bleeding events, which is currently a clinical challenge for existing commercial hemostatic solutions (e.g., argon plasma coagulation and Hemospray^R^) (*1, 36*).

Treatment of IBD can be challenging due to the limited duration of action of orally administrated drugs, which also cause systemic side effects (*37*). For example, budesonide is clinically used to treat mild to moderate active IBD including Crohn’s disease and ulcerative colitis. The duration of action for budesonide, however, is limited to 12 hours. Drug depots that can remain resident in the area of the inflammatory mucosal tissue could effectively maximize drug on target effects while minimize systemic side effects (*37*). To address this spatial-temporal challenge, we developed an e-GLUE-based drug delivery system that can be attached to specified GI mucosa for sustained therapeutics (**Fig. 4F**). For demonstration, budesonide-loaded PLGA films with a drug mass concentration of 33% as slow-release drug depots were fabricated for *in vivo* sustained therapeutic purposes due to their high drug payload (**fig. S18A**) and mechanical flexibility **(fig. S18B**). After the enema procedure, PLGA drug film was attached to the colon mucosa with the support of e-GLUE upon electrical stimulation. To mitigate the temperature rise during electrical stimulation, we applied a pulse stimulation with parameters of 15 V, 0.1 Hz in the duty cycle of 15% for four minutes (**fig. S13A**), providing additional protection to the sensitive colon tissue (**Movie S5**). e-GLUE extended the retention time of drug depots on the colon mucosa for more than four days (**Fig. 4B** and **G, Movie S6**), despite the rapid epithelium cellular renewal and mechanical shear force stemming from daily bowel movements. The drug concentration in local colon mucosa with e-GLUE treatment was significantly higher than that of oral capsule administration (**Fig. 4H**) from day 1 to day 4. Negligible drug content detected in the serum over the course of e-GLUE treatment **(fig. S18C**) indicated less systemic toxicity compared to the oral administration. No tissue damage was observed after eight days **(fig. S18D, Movie S7)**, further confirming the safety of e-GLUE technology. These results indicate that e-GLUE holds great potential as a long-term localized therapeutic platform for the treatment of chronic GI disorders in specified regions.

Sensors capable of tracking physiologic signals (e.g., tissue impedance) at defined regions of the GI mucosa can provide valuable health information, for example bowel inflammation (*30*). However, dynamic GI events, such as peristaltic waves, pose challenges for sensors to achieve stable contact and retention on the mucosa, thereby limiting measurement precision. Given its strong mucosal adhesion, we hypothesized that e-GLUE could enhance sensing precision by facilitating a robust interface between GI mucosa and sensors **(Fig. 4I)**. To validate this hypothesis, we first adhered an impedance sensor to intestinal mucosa using e-GLUE during *in vivo* tests (**fig. S19)**. e-GLUE significantly improved the contact between the rigid impedance sensor and intestinal mucosa, even during peristaltic activities. This contact improvement resulted in more precise impedance values during multiple tests (n=3, e-GLUE+ in **Fig. 4J** and **K**). Particularly, e-GLUE led to lower relative errors for both amplitude (0.3-4%) and phase (0.07-1.88%) across frequencies from 100 Hz to 100 kHz (e-GLUE+ in **Fig. 4L**), in comparison to sensors without e-GLUE support (e-GLUE-in **Fig. 4L**). Moreover, the robust mucosal retention of sensors enabled a realtime response of localized GI perturbations (e.g., temperature, **fig. S20**). Thus, e-GLUE-based sensors are able to precisely respond to localized GI environment changes through robust sensor-mucosa interfacing. Together with e-GLUE’s long-term retention (**Fig. 1E** and **Fig. 4B**), this system could provide a new paradigm for the intimate and chronic monitoring of physiological signals such as pressure and biomarkers in defined regions throughout the GI tract in awake animals.

We report the development of e-GLUE, an electroadhesive hydrogel interface that enables robust and long-lasting mucosal retention. Potential applications demonstrated mucosal hemostasis in complex GI environments, long-term localized drug delivery, and intimate biosensing applications throughout the GI tract. This system could enable clinical management of various health conditions, including gastrointestinal bleeding, inflammatory bowel disease, and monitoring of inflammation in the GI tract and beyond.

## Acknowledgment

This work was funded in part by the Novo Nordisk grant (R.L., G.T.); the Karl van Tassel (1925) Career Development Professorship and the Department of Mechanical Engineering, MIT (G.T.); Banting Postdoctoral Fellowship, NSERC Postdoctoral Fellowship (B.Y.); and MIT UROP program (H.R., S.Q., K.J.);ETH Zurich SEMP Scholarship, ZENO KARL SCHINDLER Foundation Master Thesis Grant (Y.C.); Ningbo Natural Science Foundation (Grant No. 2018A610073, Q.Z.). We thank Dr. Gary Liu, Dr. Vivian Feig, and Dr. Miguel Jimenez from MIT for insightful discussions on the manuscript; Dr. Yiyuan Yang, Dr. Ziliang Kang, Dr. Sean You, Dr. Jan-Georg Rosenboom, Dr. Ameya Kirtane, Louis DeRidder, Injoo Moon, Xisha Huang, Simon Qu, Kiwan Wong, and Koch Institute Swanson Biotechnology Center histology and imaging cores from MIT for the kind discussion and technical support; Dr. R. Bronson for help with pathology.

## Author Contributions

B.Y. conceived and designed the e-GLUE. B.Y., T.K., H.R., S.Q., S.W., K.J., Y.C., G.B., A.L., and J.L. fabricated the devices and performed the material, mechanical, and electronic characterization. B.Y. and K.N. performed the *in vivo* experiments with the assistance of J.J., A.P., J.K., K.I., N.F., and A.H. Q.Z. built the COMSOL numerical models. G.T. and R.L. provided the funding and supervised the project. All authors contributed to the writing of the manuscript.

## Data And Materials Availability

All data associated with this study are presented in the manuscript or Supplementary Materials.

## Support Information

### Methods

#### Materials

Ionic cross-linker calcium sulfate (CaSO_4_, 255548), Alginate (A2033), acrylamide (AAm, A8887), covalent cross-linker N, N′-methylenebis(acrylamide) (MBAA, M7279), free-radical initiator ammonium persulfate (APS, A3678), polymerization accelerator tetramethyl-ethylenediamine (TEMED, T7024), photoinitiator IRGACURE 2959 (I2959), acetic acid (695092), bridging polymer chitosan of medium molecular (448877), the coupling reagents, 1-ethyl-3-(3-dimethylaminopropyl) carbodiimide hydrochloride (EDC, E1769) and n-hydroxysulfosuccinimide (sulfo-NHS, 56485), Poly(diallyldimethylammonium chloride) solution (PDAC, 409022), sodium tripolyphosphate (TPP, 238503) were purchased from Sigma. Chitosan (98.2%) was purchased from CHITYOLYTIC. For all ex vivo studies, gastric tissue collections were performed within 10 min of euthanasia and the tissues were maintained in Krebs buffer and stored at 4 °C fridge during use.

#### Fabrications of e-GLUE

The synthesis of e-GLUE was prepared as follows: Briefly, chitosan powder and acrylamide were first dissolved in acetic acid solution (pH = ∼5) at 2 wt. % and 12 wt. % respectively, and stirred overnight until a clean solution was obtained. After degassing, this 5 mL of the precursor solution was mixed with 36 μL of 2 wt. % N, N′-Methylenebisacrylamide (MBAA), and 8 μL of tetramethylethylenediamine (TEMED) in one syringe (BD, 10mL). 5 mL of 20 wt. % PDAC, 226 μL of 0.27 M ammonium persulfate (APS), and 52.5 μL of 0.75M Tripolyphosphate polyanion (TPP) were injected into another syringe (BD, 10 mL). All bubbles were removed before further cross-linking. After connecting by a female luer x female luer adapter (Cole-Parmer), solutions in two syringes were mixed by pushing the syringe pistons forward and back 30 times. The mixture was stored inside a closed glass mold at room temperature overnight to allow complete polymerization. Single-network e-GLUE was prepared without Chitosan.

#### Fabrications of alginate-PAAm tough hydrogel

The synthesis of alginate-PAAm tough hydrogel was prepared following a modified protocol based on the previous report (*38*). Briefly, sodium alginate powder and acrylamide were first dissolved in distilled water at 2 wt. % and 12 wt. % respectively, and stirred overnight until a clean solution was obtained. After degassing, this 10 mL of the precursor solution was mixed with 36 μL of 2 wt. % MBAA and 8 μL of TEMED in one syringe (BD, 20mL). 226 μL of 0.27 M APS and 191 μL of 0.75M CaSO_4_ slurries were injected into another syringe (BD, 20 mL). All bubbles were removed before further cross-linking. After connecting by a female luer x female luer adapter (Cole-Parmer), solutions in two syringes were mixed by pushing the syringe pistons forward and back 10 times. The mixture was stored inside a closed glass mold at room temperature overnight to allow complete polymerization.

#### Fabrication of regular hydrogel tissue adhesives

The synthesis of hydrogel tissue adhesives was prepared following a modified protocol based on the previous report (*31*). Briefly, the alginate-PAAm tough hydrogel surface was treated with a mixture of bridge polymer and coupling reagents for the carbodiimide coupling reaction. The bridge polymer chitosan was dissolved into distilled water at 2.0 wt. % and the pH was adjusted to 5.5∼6 by acetic acid. EDC and NHS were used as the coupling reagents. The final concentrations of EDC and NHS in the solution of the bridging polymer were both 12 mg/mL. During GI mucosa bonding, the entire device was pressed on the mucosa for minutes using an endoscope to guarantee a sufficient reaction.

#### Polymerization of cations

Cations (3ATAC, 3METAC, 2ATAC, 2DAMA, N3DMA) were dissolved into DI water in a concentration of 10 wt.%, respectively. After adding I2959, the cations solutions were exposed to UV (256 nm) for 1 h to allow a complete polymerization.

#### e-GLUE electrode assembly fabrication

e-GLUE electrode assemblies were manually manufactured by assembling four layers: negative electrode, VHB foam, a positive electrode, and PET film. All features were designed in Solidworks (Dassault Systemes) and cut using a laser cutting machine (MIYACHI, WL-100A). Copper wires were connected to electrodes with epoxy sealed to avoid shorting issues. Rigid Heat Shrink Sleeving was used to encapsulate the copper wires to guide through the endoscopic channel.

#### Strength measurement of e-GLUE

Rectangular strips of e-GLUE (80 × 25 × 4 mm^3^) were glued to two rigid acrylate clamps (80 × 10 × 1.5 mm3). Samples were prepared for pure shear tests with a universal testing machine (Instron, Model 5543, the loading cell is 250N). The tensile strain rate was fixed at 200% min^−1^. The tensile strain (ε) was defined as the length change (Δl) divided by the original length (l_0_) of the sample. *Young’s* modulus was calculated from the linear region of the strain-stress curve.

#### Adhesion performance measurement

Adhesion performance was measured with either 180°-peeling or tensile tests. 180°-peeling test: Under electrical stimulation, a ribbon of the e-GLUE (80 × 15 × 1.4 mm^3^) was adhered to a mucosa mimic gel (i.e., Alg-PAAm hydrogel) with one end open, forming a bilayer with an edge crack. The surfaces of e-GLUE and mucosa mimic gel were bonded to a rigid polyethylene terephthalate (PET) film with super glue, to limit deformation to the crack tip. The free ends of e-GLUE and gel were attached to plastic sheets, to which the machine grips were attached. The loading rate was set constant at 100 mm min-1. Adhesion energy, namely the energy required to increase a unit area of the e-GLUE-substrate interfacial crack. The adhesion energy was two times the plateau value of the ratio of the force and width. Both force and displacement were recorded continuously throughout the experiment. A minimum of three specimens were used for all mechanical test conditions. Tensile test. Due to the unique configuration of the e-GLUE-mucosa bilayer structure during non-invasive adhesion. A manual mechanical testing stage (Mark-10, Model ES20) coupled with a force gauge (Mark-10, Model M4-05) was used to apply a precisely controlled tensile pulling force on e-GLUE, which was then converted to adhesive strength using contacting area.

#### Cytocompatibility

The cytotoxicity of materials to NIH/3T3 was evaluated following standard protocols as described in International Organization for Standardization (ISO) 10993-12: Biological evaluation of medical devices--Part 12: Sample preparation and reference materials. Complete DMEM was first prepared by supplementing 1% nonessential amino acid, 1% penicillin-streptomycin, and 10% fetal bovine serum. Material extracts were prepared by adding 1 mL of complete DMEM to every 200 mg of e-GLUE in a conical tube to completely cover the hydrogels. On the same day, cells were seeded into 96-well plates with a density of 10,000 cells per well. After 24 hours of incubation at 37°C, the extracts were collected. Then, the cell culture medium within the 96-well plate was removed and replaced with the supplemented hydrogel extracts. Complete DMEM incubated at the same condition of extraction was used as a control. The treated cells were incubated at 37°C, 95% relative humidity, and 5% CO_2_ atmosphere. After 24 hours, the cell viability was determined using a LIVE/DEAD viability kit (Invitrogen, L3224) and an MTT viability test kit (Sigma-Aldrich, 11465007001), respectively, according to the manufacturer’s protocol. The LIVE/DEAD stained cells were visualized using a confocal laser scanning microscope (Zeiss, LSM710). Live cells were shown in green fluorescence and dead cells were shown in red. The MTT test was examined using a microplate reader (Synergy HTX, Agilent). The wavelength of 570 nm was used for absorbance reading while 650 nm was used as reference. Cell viability was calculated by (I_(570, sample)-I_(650,sample))/(I_(570,control)-I_(650,control) )×100%. Materials with over 70% viability were deemed cytocompatible following ISO 10993-5: Biological evaluation of medical devices. Five replicates were used for LIVE/DEAD staining while seven were for MTT assay.

#### Synthesis of FITC-labeled quaternary chitosan for diffusion visualization

A buffer solution (pH=9.5) was prepared by mixing 50 mM Na_2_CO_3_ and 50 mM NaHCO_3_ solutions accordingly. After dissolving 300 mg quaternary chitosan into 180 mL of buffer solution and a 15-minute degassing, 5.7 mL of 2 mg/mL FITC dimethylformamide (DMF) solution was added into the chitosan solution under stirring drop by drop. The mixture was stored inside a closed beaker with 12 h stirring at room temperature to allow a complete reaction. The glass beaker was covered from light. The mixture was freeze-dried for future tests after dialysis for 4-5 days.

#### Confocal microscopy

We used FITC-labeled quaternary chitosan to track the diffusion of polycation chains from e-GLUE into mucosal tissues. FITC-labeled e-GLUE was first bonded to mucosa under different stimulation conditions. The entire sample was then imaged with confocal fluorescence microscopy (Leica TCS-sp5), with an excitation wavelength of 490 nm and emission wavelength of 525 nm for FITC. A series of confocal images were taken by scanning samples along its thickness.

##### *In vivo* experiments

All animal experiments were conducted following the protocols approved by the Committee on Animal Care at the Massachusetts Institute of Technology. Swine models were chosen because of their anatomical similarity to humans in GI-related studies (*39*). Randomization of the animals was not performed. Female Yorkshire swine (Cummings Veterinary School at Tufts University in Grafton, MA) weighing 43–90 kg and aged 3–6 months were used. The swine were placed on a liquid diet 24 h before the study and fasted on the day of the procedure. On the morning of the procedure, the swine were sedated using intramuscular injection of either 5 mg kg^−1^ telazol (tiletamine/zolazepam), 2 mg kg^−1^ xylazine and 0.04 mg kg^−1^ atropine, or 0.25 mg kg^−1^ midazolam and 0.03 mg kg^−1^ dexmedetomidine. After intubation, anesthesia was maintained with isoflurane (2–3% oxygen). Under anesthesia, vital signs were monitored and recorded every 15 min throughout the study. After the study, swine was woken up using intramuscular injection of 0.1 mg kg^−1^ atipamezole and monitored closely during the recovery process until full recovery was achieved. For studies on ambulating animals to monitor long-term gastric retention of the e-GLUE, the swine was fed normally and anesthetized 1-2 times weekly for radiography.

##### *In vivo* retention and safety evaluation

e-GLUE devices were endoscopically delivered and attached to the gastric mucosa of a swine using an overtube. After electrical stimulation and adhesion, Weekly X-ray radiographs were subsequently performed to determine the residency time of the devices. X-rays were taken until all devices passed. During the device retention, the animals were evaluated clinically with no evidence of any changes in feeding or stooling patterns. In addition, the long-term safety of mucosal tissues including the duodenum, colon, and stomach treated by e-GLUE was examined both visually and through histological analysis after 7-60 days.

#### Histology

Histological analysis was performed on GI tissue biopsies to characterize the safety. Biopsies were taken from the e-GLUE-treated mucosa from the stomach, SI, and colon. The biopsies were fixed in formalin fixative (Sigma Aldrich) for 72 hours before transfer to 70% ethanol. Tissue samples were then embedded in paraffin, cut into 5-μm-thick tissue sections, stained with hematoxylin and eosin, and imaged using an Aperio AT2 Slide Scanner (Leica Biosystems). These samples were analyzed by a board-certified pathologist.

##### *In vivo* demonstrations

Specific methods for mucosal sealing, drug delivery, and intimate sensing in pigs are described within their respective sections. For drug delivery demonstration, the swine was fed normally and anesthetized 2 times weekly for localized biopsy in the colon. All serum samples were combined with acetonitrile in a 1:3 ratio (v/v) and then centrifuged at 1200 rpm at 4°C for 15 min for protein precipitation and extraction. The supernatant of each tube was then loaded into microtubes and processed using HPLC to quantify the drug concentrations.

#### Synthesis of budesonide-PLGA films as slow-release drug depots

Budesonide–PLGA patches were synthesized by uniform precipitation from organic solvents. 120 mg of budesonide and PLGA (PLGA: budesonide = 1:1, 1:1.5, 1:2) were dissolved in 5 mL acetone. The mixture was vortexed and sonicated for 0.5 h to ensure complete dissolution. Then the mixture was poured into a rectangle glass mold (25 mm*20 mm) covered with aluminum foil. After 2 days, the budesonide–PLGA films were taken off the glass mold and cut into 15 mm*3 mm patches, which have a thickness of around 200 μm.

##### *In-vitro* release of budesonide–PLGA film

The in-vitro release of budesonide from budesonide–PLGA film was performed using a horizontal shaker with 250 r.p.m. speed at 37 °C (Innova Shaker). PBS (pH 7.4, ×1, Gibco, Thermo Fisher Scientific) was selected as the *in-vitro* medium for drug release kinetics characterization given the intent to target the rectal mucosa environment where extracellular fluid with a similar pH would be encountered. Each budesonide-PLGA patch was added to 10 mL PBS with 1% Tween 20. Experiments were performed at 37 °C, and 1 mL samples were taken daily up to 14 days of release. Buffers were refreshed at different time intervals, and the drug content was analyzed using HPLC analysis.

#### High Performance Liquid Chromatography (HPLC) analysis of Budesonide release

High Performance Liquid Chromatography (HPLC) was used to determine the drug concentrations from all in-vitro release assays. An Agilent 1260 Infinity II HPLC system equipped with a quaternary pump, autosampler, thermostat, control module, and diode array detector were utilized as described previously. Data processing and analysis was performed using OpenLab CDS ChemStation®. Budesonide was separated on an Agilent Poroshell 120 EC-C18 analytical column 3.0 x 50 mm with 2.7 μm particles, maintained at 40 °C. The optimized mobile phase consisted of A: 0.1% formic acid in water and B: acetonitrile. Isocratic elution (30% A, 70% B) was employed over 5 minutes at a flow rate of 0.750 mL/min. The injection volume was 10 μL, and the selected ultraviolet (UV) detection wavelength was 244 nm at a bandwidth of 4.0, no reference wavelength, and an acquisition rate of 10 Hz.

#### Liquid Chromatography Triple Quadrupole Mass Spectrometry (LC-MS/MS) - Serum/Plasma and Tissue Biopsy Analysis

Plasma samples were prepared using protein precipitation in acetonitrile in a 1:3 mixture. Specifically, 100 uL of serum or plasma was spiked with 50 uL of either spiking solution (used to generate calibration curve – budesonide standards pre-dissolved in 100% acetonitrile) or 100% acetonitrile for measured samples. After pipette mixing, 250 uL of extraction solution (200 ppb ISTD mixture in 100% acetonitrile) is added to this mixture, and the sample is vortexed for 15 seconds. After pelleting the sample at 21,000 g for 10 minutes at 4 °C, the supernatant is transferred to a micronic vessel for analysis.

Tissue samples are prepared in a similar manner. The tissue biopsy is weighed in an omni scientific homogenization bead mill tube. After the mass is recorded, 50 uL of either spiking solution (used to generate calibration curve – budesonide standards pre-dissolved in 100% acetonitrile) or 100% acetonitrile for measured samples. Subsequently, 250 uL of extraction solution (200 ppb ISTD mixture in 100% acetonitrile) is added to this mixture and the tube is homogenized for 2 minutes. The sample is then pelleted at 21,000 g for 10 minutes at 4 °C, and the supernatant is transferred to a micronic vessel for analysis.

Liquid Chromatography Triple Quadrupole Mass Spectrometry (LQ-MS/MS) was used to determine model drug concentrations in all in-vivo serum and plasma samples. An Agilent 1290 UHPLC system with a binary pump, autosampler, and thermostat was coupled to an Agilent 6495B triple quadrupole mass spectrometer. Data sets were generated using the Masshunter® LC-MS control suite, and data processing and analysis was performed in QuantMyWay®.

Budesonide and Budesonide-d8 were separated on an Agilent Poroshell 120 EC-C18 analytical column 3.0 x 50 mm with 2.7 μm particles, maintained at 40 °C. The optimized mobile phase consisted of A: 0.1% formic acid in water and B: acetonitrile. Isocratic elution (30% A, 70% B) was employed over 5 minutes at a flow rate of 0.750 mL/min. The injection volume was 10 μL. The compounds underwent electrospray ionization (positive mode) with drying gas temperature of 250 °C, sheath gas temperature of 380 °C, sheath gas flow rate of 10 L/min, and drying as flow of 16 L/min. The nebulizer was kept at 35 psi. The capillary voltage was set to 4000 V in positive mode, and there was no in-source fragmentation. Budesonide and budesonide-d8 were monitored under dynamic MRM (multiple reaction monitoring) with transitions of 431.3 m/z → 323.2 m/z and 439.3 m/z → 323.2 m/z for budesonide and budesonide-d8, respectively. The collision energies were set as 12 V and 20 V for naloxone and naltrexone, respectively. An LLOQ of 0.100 ng/mL was achieved using the sample preparatory and analytical methodologies conveyed in this report.

#### Statistical analysis

Statistical analyses were performed on GraphPad Prism software (GraphPad Software, Inc.). Results are depicted as mean ± standard deviation (SD). We conducted a two-sided student’s t-test to analyze the statistical differences in experiment results. *P* values < 0.05 were considered statistically significant.

