## Supplemental Information for "Electroadhesive hydrogel interface for prolonged mucosal theranostics"

**Table of contents – Supplementary Information**

**Supplementary Text**

**Figure S1-S20**

**Movie S1-7**

**Supplementary Text**

**Abbreviation**

PDAC, poly(diallyldimethylammonium chloride)

P3ATAC, poly[(3-Acrylamidopropyl) trimethylammonium chloride]

P3METAC, poly{[3-(Methacryloylamino) propyl]trimethylammonium chloride}

P2ATAC, poly {[2-(Acryloyloxy)ethyl] trimethylammonium chloride}

P2DAMA, poly{2-(Dimethylamino)ethyl methacrylate}

PN3DMA, poly{N-[3-(Dimethylamino)propyl]methacrylamide}

**Numeric simulations**

All 3D models and simulations were performed using COMSOL Multiphysics (version 6.0). The physical model incorporates the electrode current (ec), bioheat, and multiphasic modulus of COMSOL. The electrode current modulus simulates the electrical field, potential voltage, and current density. The bioheat modulus simulates heat transfer in tissue and gel, while the multiphasic modulus accounts for the heat generated by the electric field as a heat source in the bioheat modulus.

It's worth noting that although the electrical conductivity of tissue and gel can change with temperature, our model simplifies this aspect. These materials typically exhibit less than a 10% variation in electrical conductivity unless the temperature exceeds their ignition point, resulting in a dramatic decrease in electrical conductivity.

The term 'Tetrahedral Mesh' refers to a type of mesh formed by dividing objects into tetrahedrons in three-dimensional space. In the field of biomedical engineering, the tetrahedral mesh is commonly used for modeling and simulating human organs.

The electromagnetic and heat transfer parameters for mucosal tissues were derived from numerical values associated with animal muscle. Specifically, the electrical conductivity of these tissues was set at 0.5 S/m, while the heat conduction was assigned a value of 0.49 W/(mK). Additionally, the heat capacity was determined to be 3421 J/(kg*K), and these tissues were considered isotropic in their properties.

In contrast, the electrical conductivity of the gel used in our experiments was obtained from experimental data and found to be 1.5 S/m. The heat transfer parameters of the gel were assumed to be equivalent to those of water.

The simulations involved the instantaneous establishment of an electrical field upon the application of electrical potential to the tissue. As a result, a static simulation model was utilized, while the temperature within the tissue and gel evolved over time due to the generation of Joule heat.

**Heat transfer**

"The Pennes’ equation was employed to describe heat transfer within the tissue:

$$\rho_{t}c_{t}\frac{\partial T_{t}}{\partial t}=\nabla k_{t}\nabla T_{t}+\rho_{b}\omega_{b}\left( T_{b}-T_{t} \right)+Q_{meta}+Q_{SAR}$$

Where 𝜌*_t_*, 𝑐*_t_*, 𝑘*_t_*, *T_t_* represent the density, heat capacity, heat conductivity, and temperature of the tissue, respectively. Similarly, 𝜌*_b_ ,T_b_*, *w_b_* represent the density, temperature, and blood perfusion of the tissue. Q_SAR_, Q_meta_ denote the volumetric heating rate of Radiofrequency and tissue metabolic heating rate.

For the boundaries of the tissue, electrode, and ground pad, they were in contact with either air or cooling water, and heat exchange occurred through heat convection:

$$k\nabla T|_{bourdary}=h_{s}(T_{air}-T_{t})$$

Where *h_s_* represents the natural heat convection coefficient of passive airflow, and *T_air_* is the temperature of the surrounding air.

**Electrode field**

Assuming current can flow in all fields of model, the static electrical field in the tissue can be applied by Maxwell equation.

$$\nabla\cdot\left( -\sigma_{t}\nabla V \right)=0$$

Where *V* and 𝜎*_t_* presented the electric potential and electric conductivity of tissue.

The heat generated by Radiofrequency can be calculated by the electric potential gradient and electric conductivity of tissue.

$$Q_{SAR}=\sigma_{t}{(\nabla V)}^{2}$$

The outer surface is surrounded by air, whose electric conductivity is 10^-15^ S/m. Therefore, the outer surface could be considered as insulation.

$$\nabla V=0$$

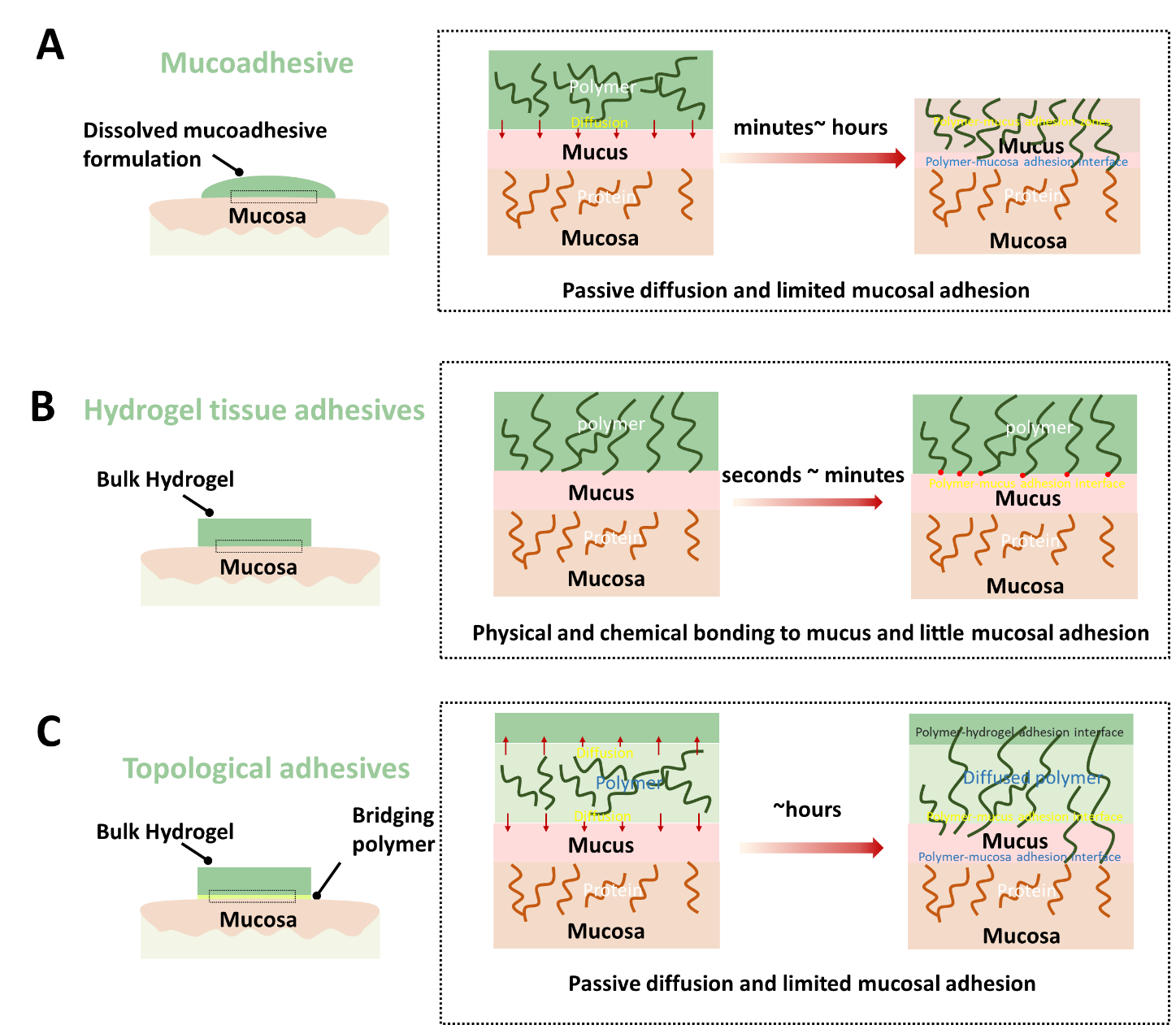

**fig. S1: Exiting adhesion mechanisms for GI mucosa bonding. A)** mucoadhesive; **B)** hydrogel tissue adhesives; **C)** topological adhesives.

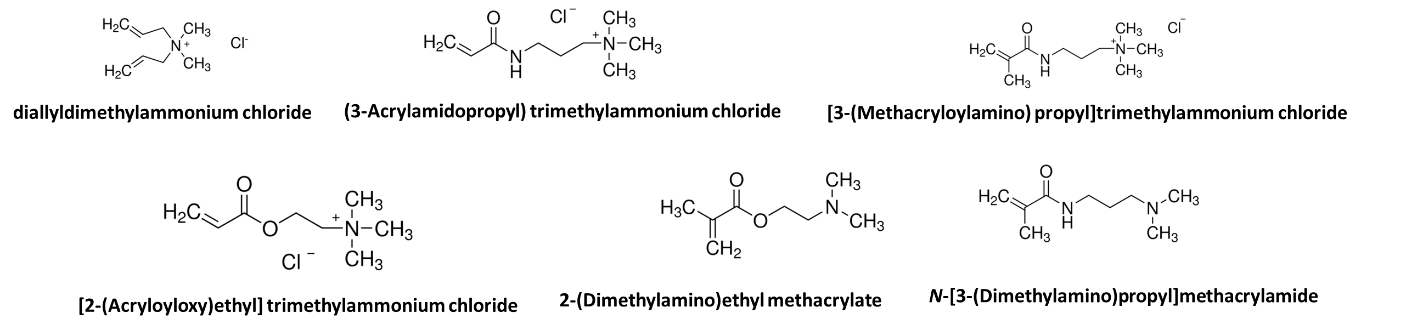

**fig. S2. Chemical structures of various cationic monomers:** diallyldimethylammonium chloride, (3-Acrylamidopropyl) trimethylammonium chloride, [3-(Methacryloylamino) propyl]trimethylammonium chloride, [2-(Acryloyloxy)ethyl] trimethylammonium chloride, 2-(Dimethylamino)ethyl methacrylate, N-[3-(Dimethylamino)propyl]methacrylamide.

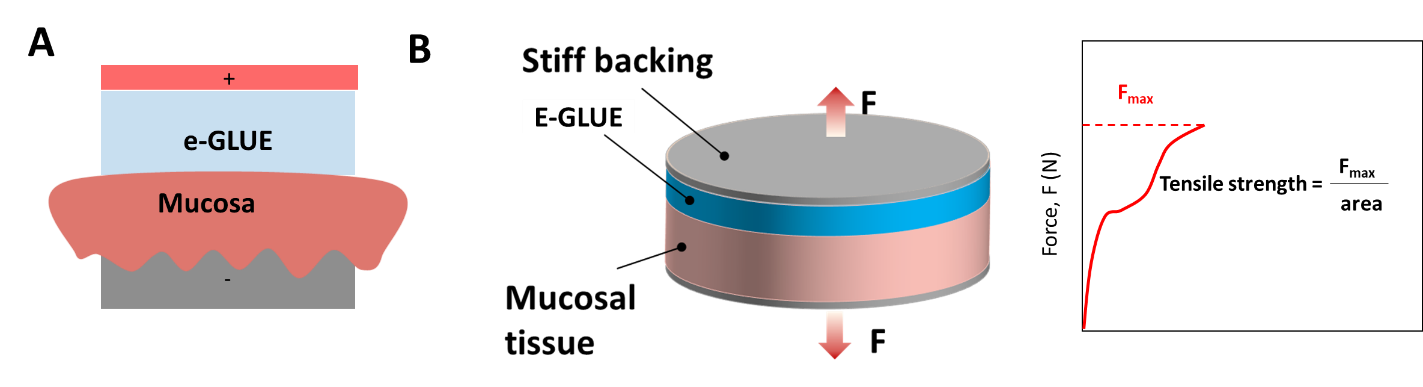

**fig. S3. Electroadhesion and their adhesion characterization.** **A)** Forming adhesion between e-GLUE and mucosa through a sandwiched electrode configuration (invasive)**. B)** Tensile test setup to characterize e-GLUE’s adhesion strength on mucosa. The tensile strength is determined by F_max_/ sectional area.

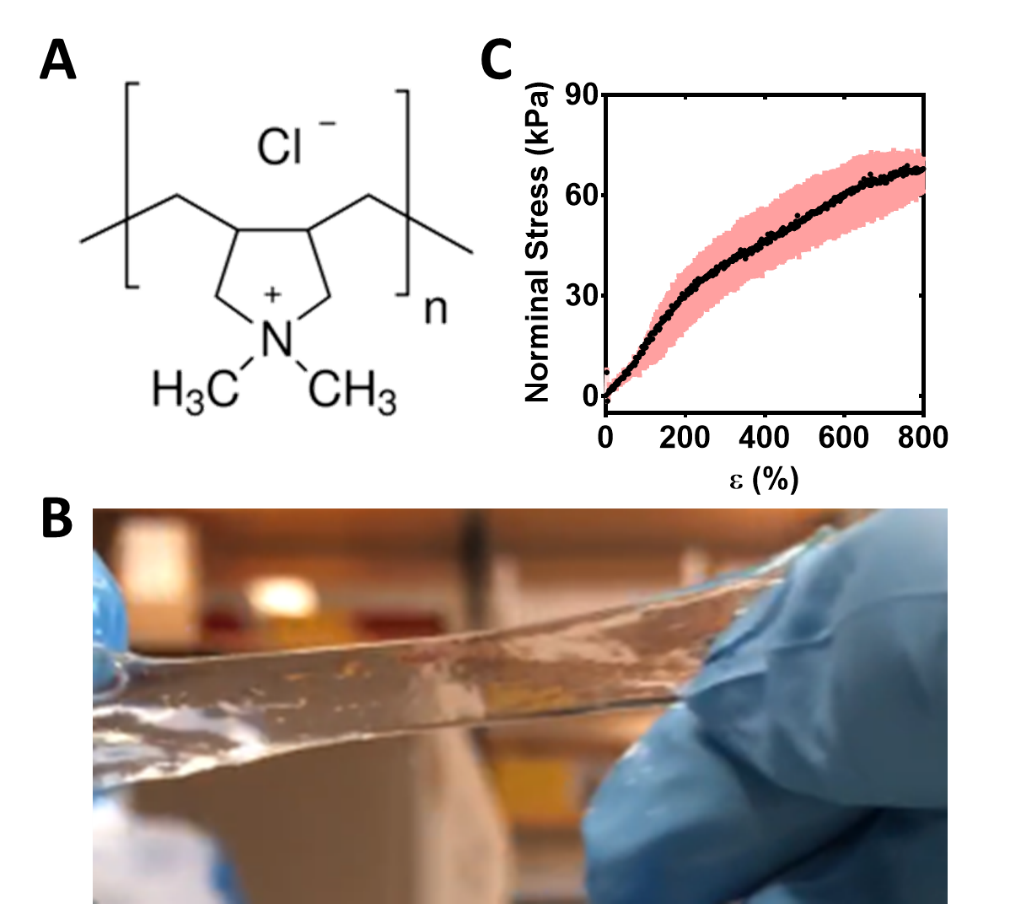

**fig. S4: PDAC-based e-GLUE. A)** Chemical structure of PDAC. **B)** Mechanical stretchability and **C)** representative strain-stress curve of PDAC-based chitosan-PAAm double-network hydrogel (PDAC-based e-GLUE). The values in panel C indicate the mean and the standard deviation (n = 3).

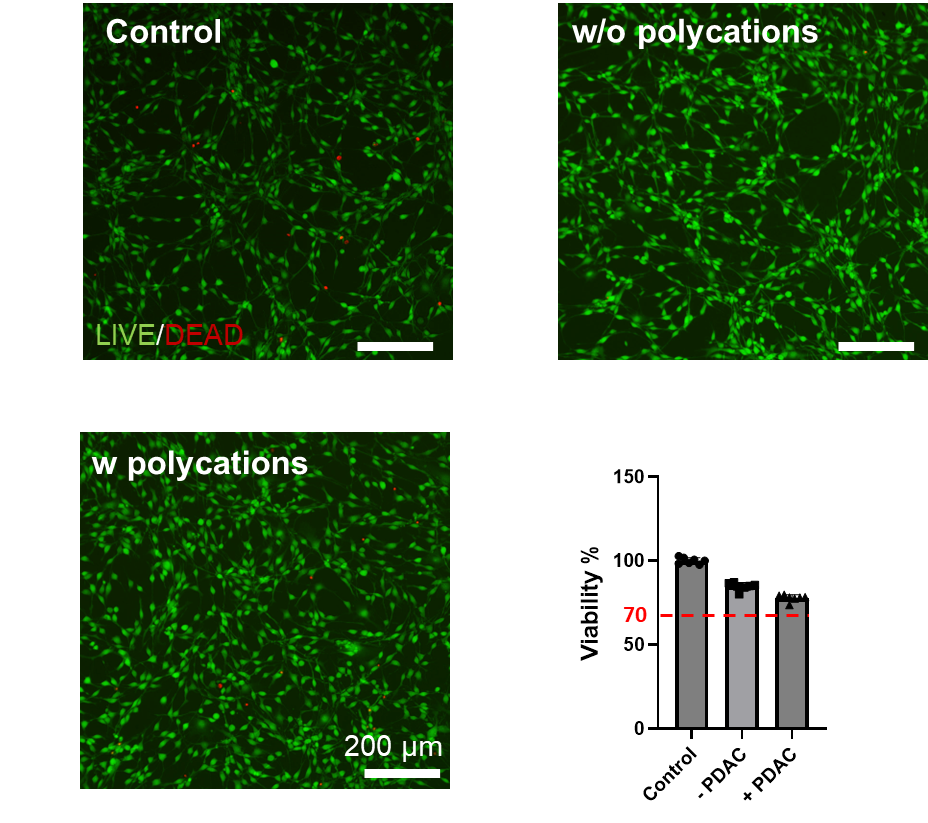

**fig. S5. In vitro biocompatibility of the e-GLUE in a live/dead assay of mouse embryonic fibroblast cells (NIH/3T3) after 24 hours of culture.** MTT viability assay indicated that cells maintained 78% ± 2% (n=7) of their metabolic activity compared to the negative control, exceeding the 70% safety threshold suggested by the ISO standard. Negative control: cell culture on a standard 96-well plate. The test follows the International Organization for Standardization (ISO) 109003-5: Tests for in vitro cytotoxicity standard.

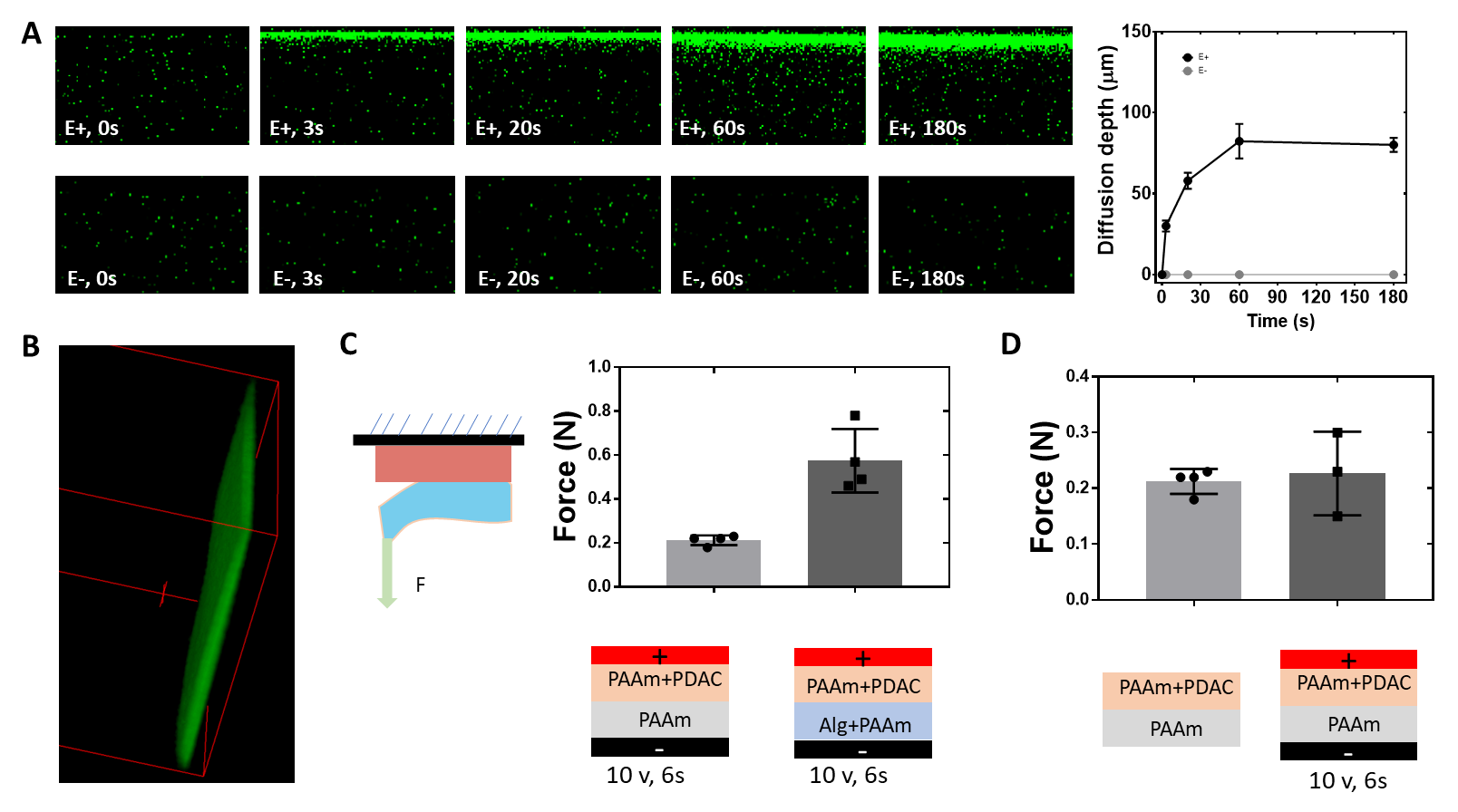

**fig. S6. Investigation of electroadhesion mechanism between the e-GLUE and mucosa. (A)** Active diffusion of FITC-conjugated quaternary chitosan into the mucosa under different time points, with (*E+*) and without (*E-*) electrical stimulation (U = 3V, the thickness of FITC-conjugated e-GLUE is 0.75 mm). Diffusion depth increased with the stimulation time before reaching to its plateau. Diffusion of FITC-chitosan was imaged by confocal fluorescence microscopy. The values indicate the mean and the standard deviation (n = 6). (**B**) 3D view of the diffusion depth into mucosal layers after 3-min electrical stimulation. (**C**) Adhesion force of e-GLUE and PAAm hydrogel, e-GLUE and Alg-PAAm hydrogel under electrical stimulation (10 V, 6 s). 90^o^-peeling test was conducted to measure the adhesion force. (**D**) Adhesion force of e-GLUE and PAAm hydrogel with and without electrical stimulation (10 V, 6 s). Such an electrical condition was chosen to eliminate the significant temperature rise. The values in panels **C** and **D** indicate the mean and the standard deviation (n = 3–4).

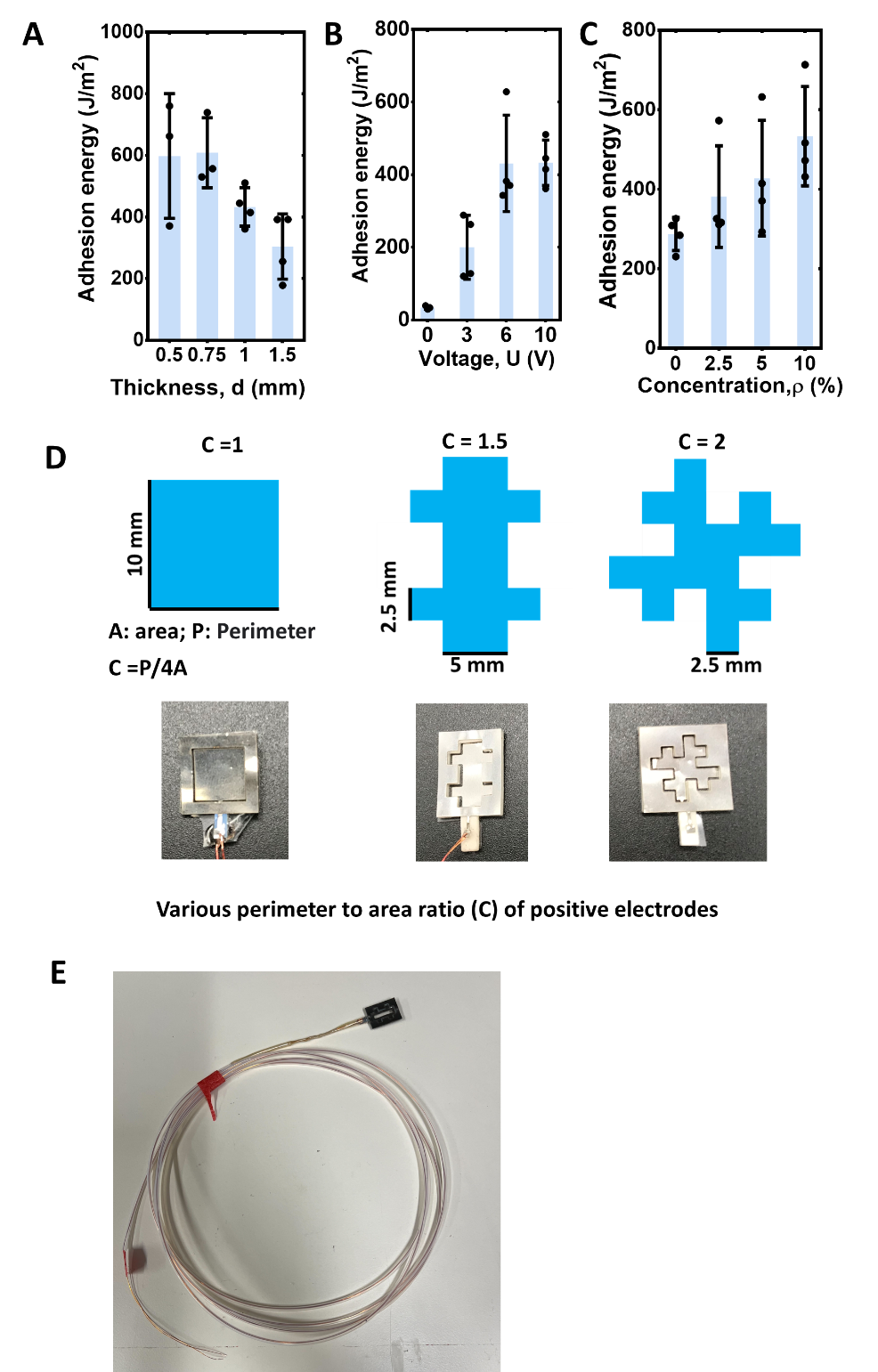

**fig. S7. Optimization of e-GLUE and its electrode** **assembly.** The adhesion energy of e-GLUE depends on **A**) the gel thickness (d), **B**) the voltage amplitude (U), and **C**) the polycation concentration (ρ). **D**) e-GLUE electrode optimization under three perimeter-to-area ratios (C:1, 1.5, and 2) and their optical photo after assembling. **E**) A photo of home-made e-GLUE electrode assembly (C= 1.5) before the insertion into the endoscope channel. Condition used for (**A**): U=10 V, t=10 s, ρ=10 %, and Mw=200~350 K; for (**B**): d= 1 mm, t=10 s, ρ=10 %, and Mw=200~350 K; for (**C**): d= 1 mm, U=10 V, t=3 s, ρ=10 %, and Mw=200~350 K. The values in panels **A, B** and **C** indicate the mean and the standard deviation (n = 3–4).

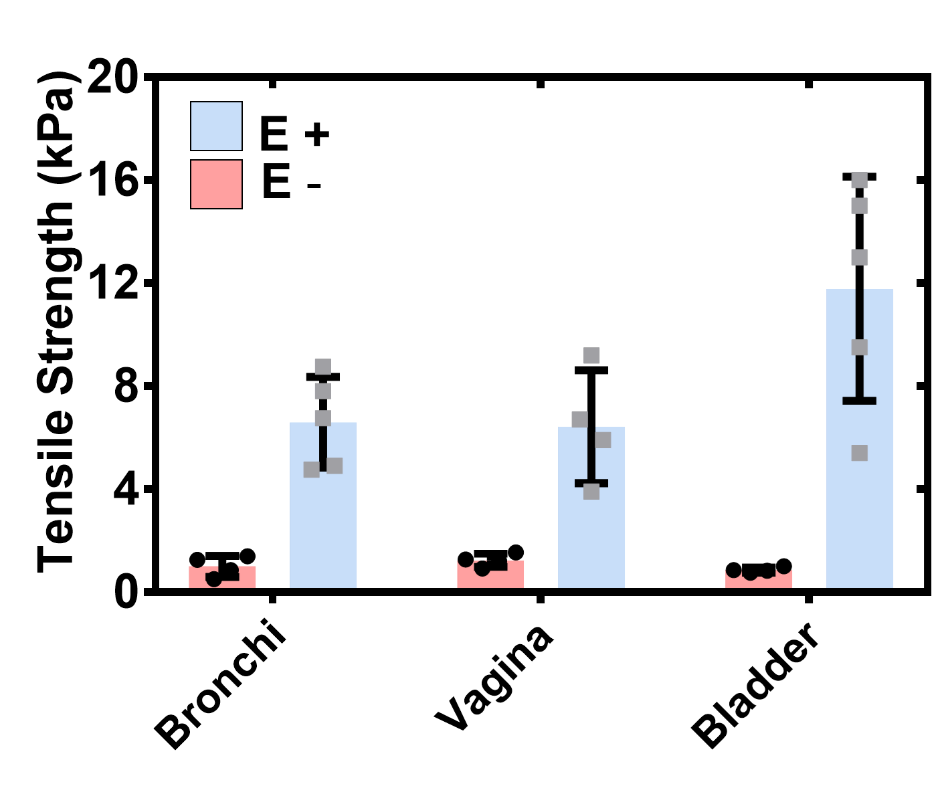

**fig. S8.** Tensile strength of bronchi, vagina, and bladder mucosal tissues adhered by e-GLUE with **(E+**) and without (**E-**) electrical stimulation (15 V, 40 s). Square-shaped electrodes were used here for non-invasive bonding. e-GLUE thickness is 0.75 mm and polycations concentration is 10%. The values indicate the mean and the standard deviation (n = 4–5).

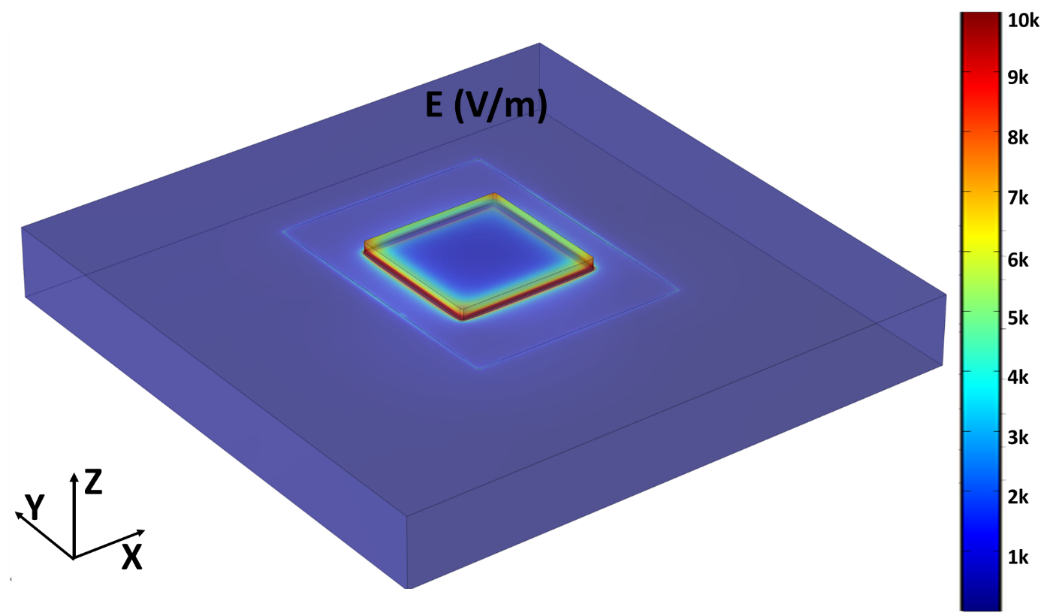

**fig. S9.** Numerical modeling of the electrical field distribution of novel e-GLUE electrode design (Square shape, C= 1). The voltage amplitude was set at 10 V.

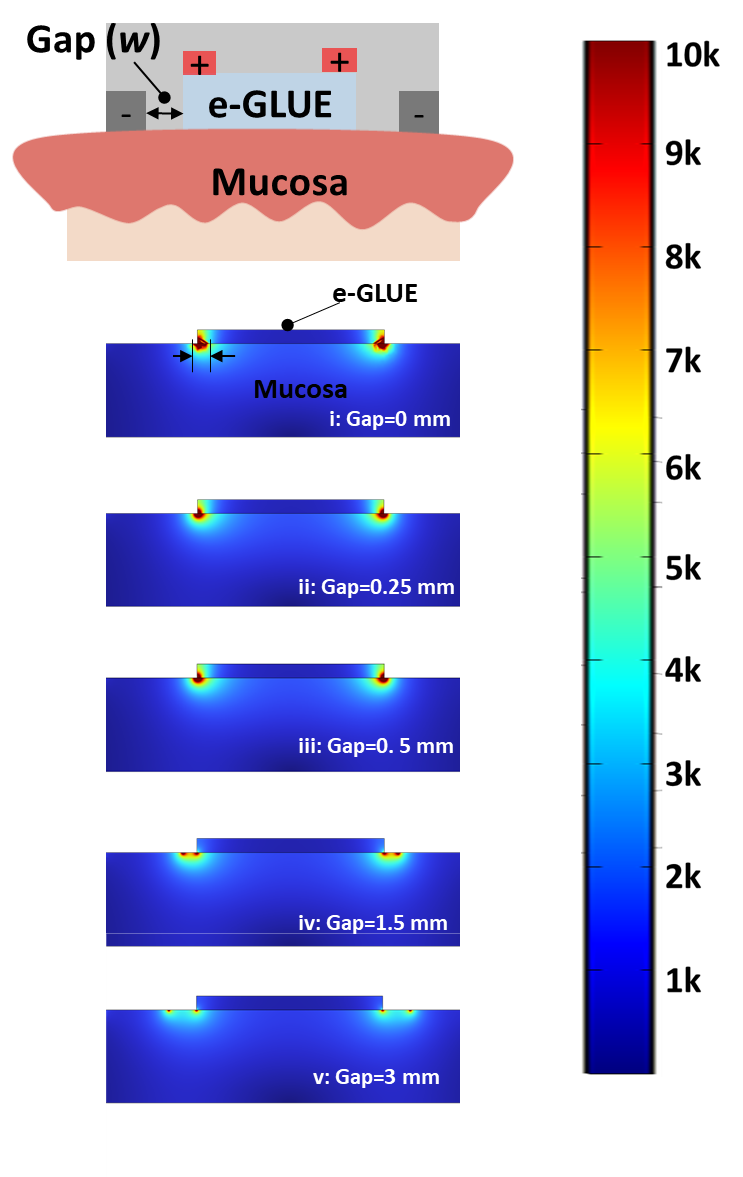

**fig. S10.** The change of electrical field distribution under various gaps between positive and negative electrodes (C= 1.5). The voltage amplitude was set at 10 V.

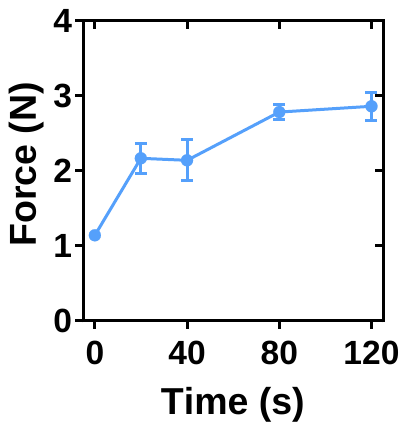

**fig. S11.** Adhesion force of SI mucosal tissues adhered by e-GLUE under different stimulation time. Non-invasive electrodes (*C=1.5*) were used here for tissue bonding. e-GLUE thickness is 0.75 mm and polycations concentration is 10%. The voltage amplitude was set at 10 V. The values indicate the mean and the standard deviation (n = 4).

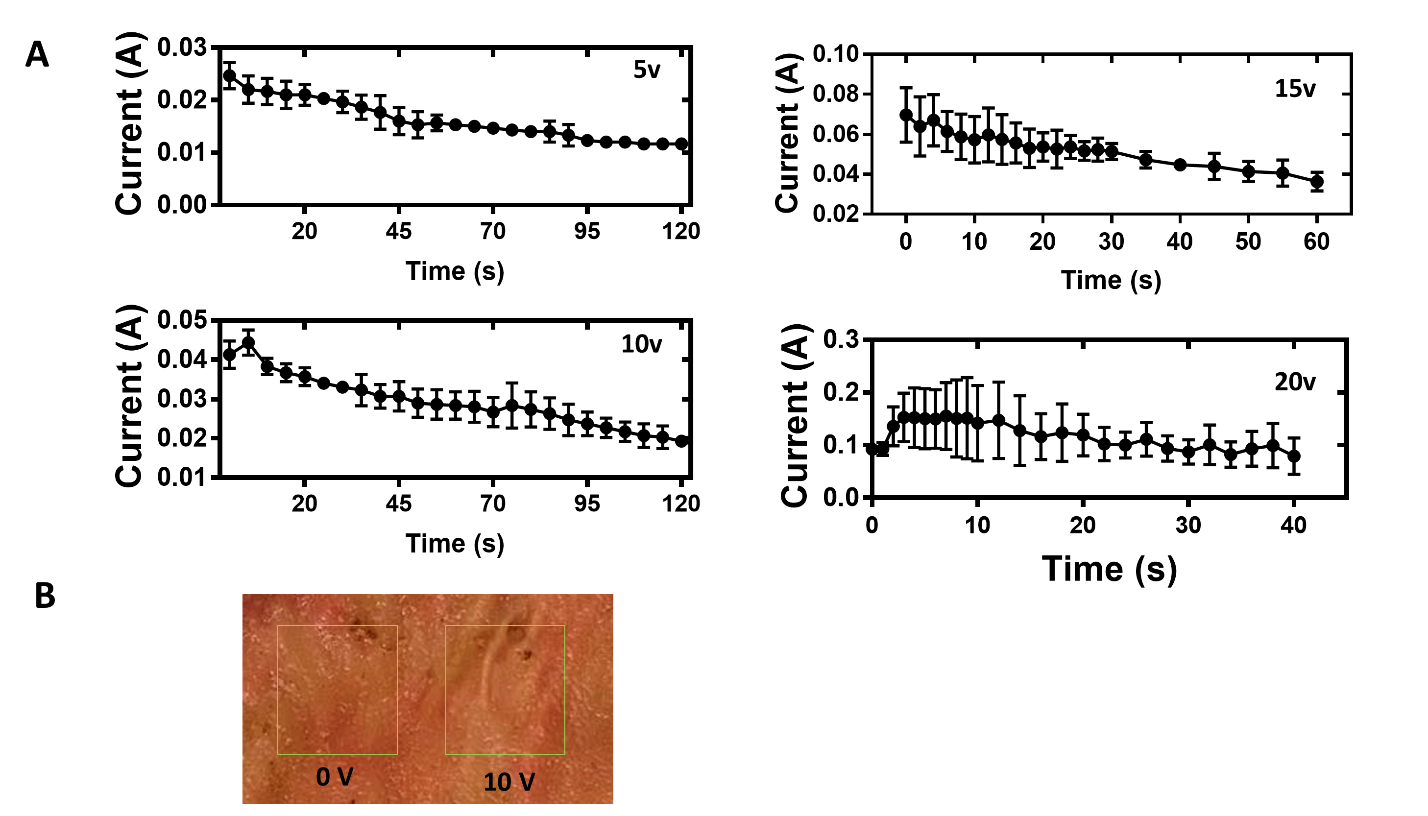

**fig. S12. Current and tissue damage monitoring under different stimulation conditions**. **A)** Current monitoring under different stimulation voltages. Non-invasive electrodes (*C=1.5*) were used here for SI mucosal bonding. e-GLUE thickness is 0.75 mm and polycations concentration is 10%. **B**) Tissue damage observations on the *ex-vivo* SI after 80 s. The values in panel **A** indicate the mean and the standard deviation (n = 3).

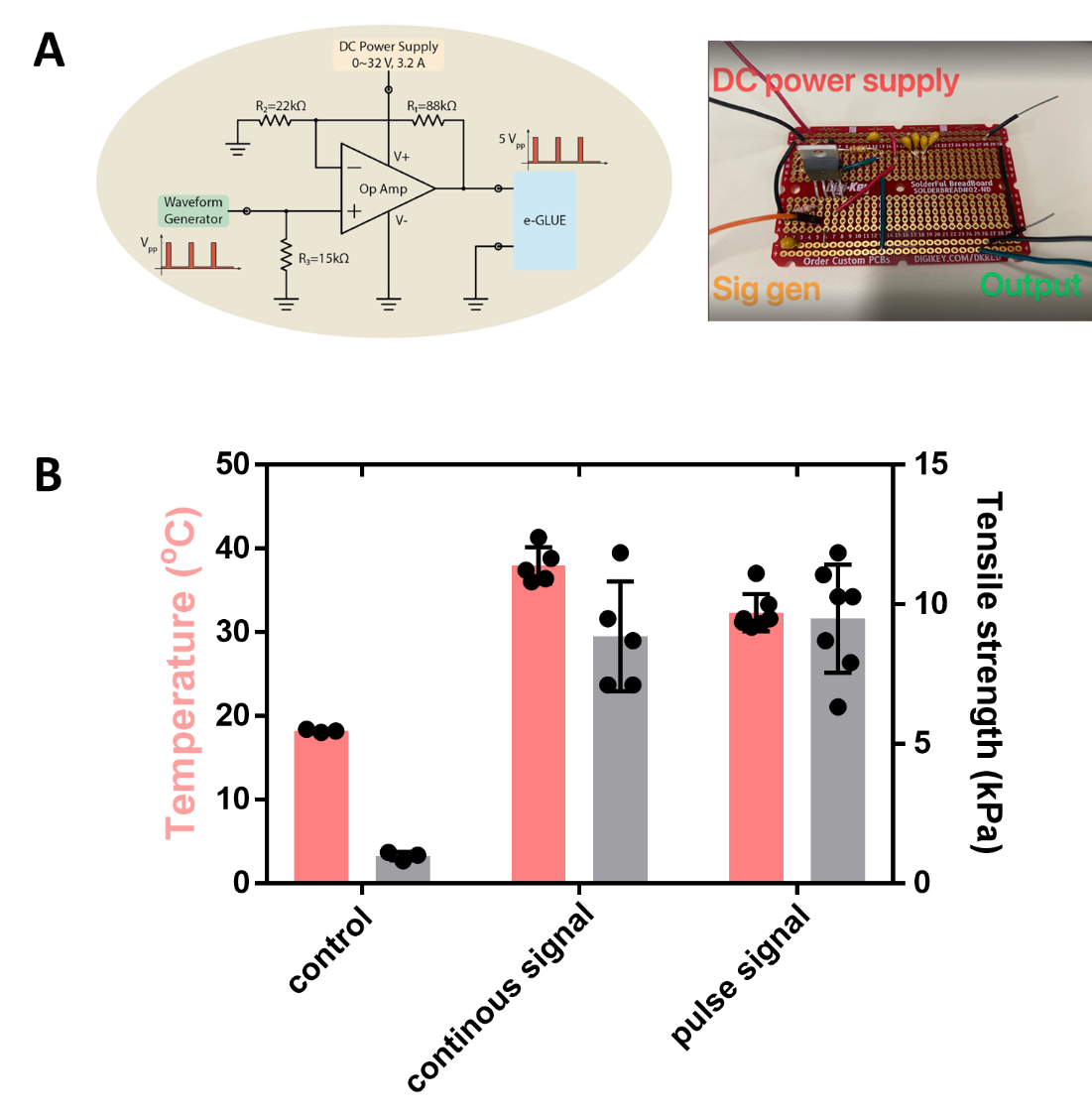

**fig. S13. Temperature rise elimination through pulse stimulation.** **A**) Circuit diagram and PCB for pulse signal generation. **B**) The comparison of temperature and adhesion strength under different signal types. Continuous signal was set at 15V for 40 s. Pulse signal was set at 15V for 200s with the frequency of 0.1 Hz and duty cycle of 20%. The values in panel **B** indicate the mean and the standard deviation (n = 3–7).

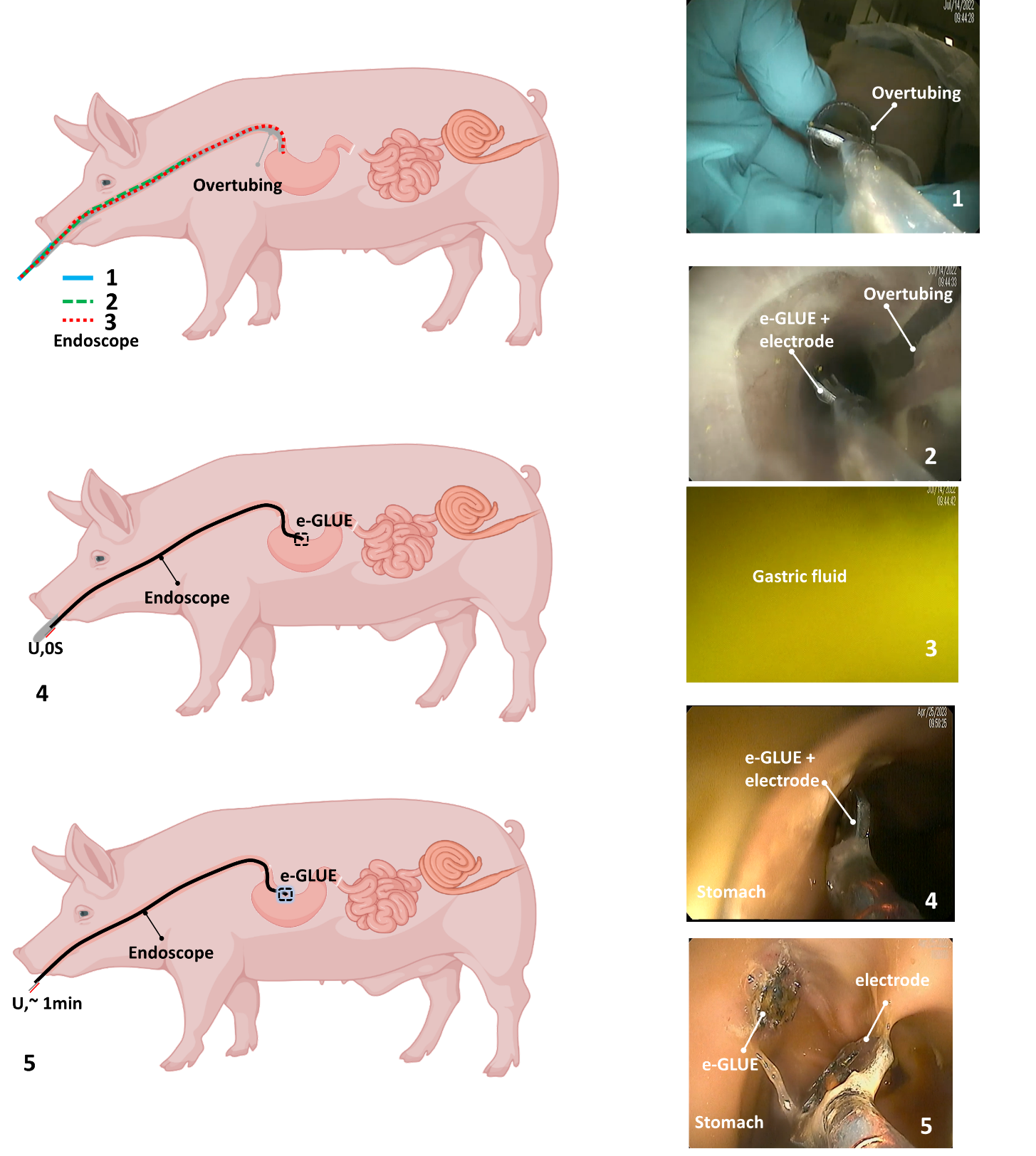

**fig. S14.** e-GLUE delivery procedures for retention performance validation in the stomach *in vivo*.

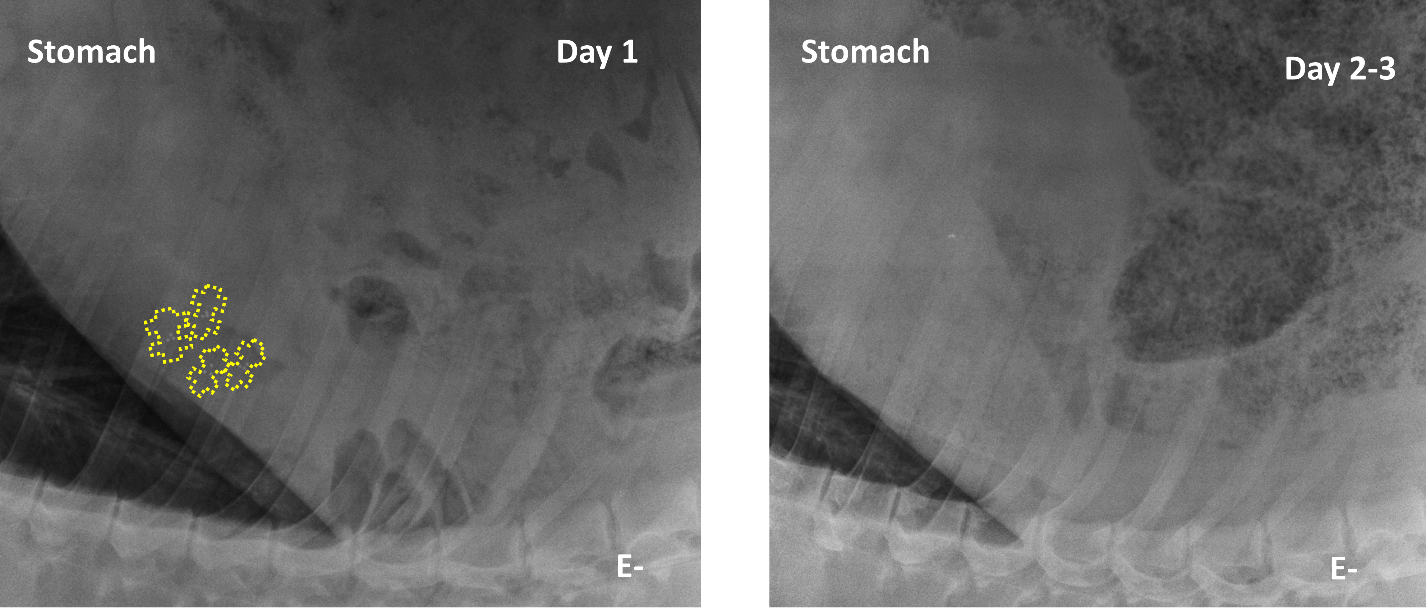

**fig. S15.** X-ray images show that the retention time of regular hydrogel adhesives on the gastric mucosa is less than 2 days, n=4.

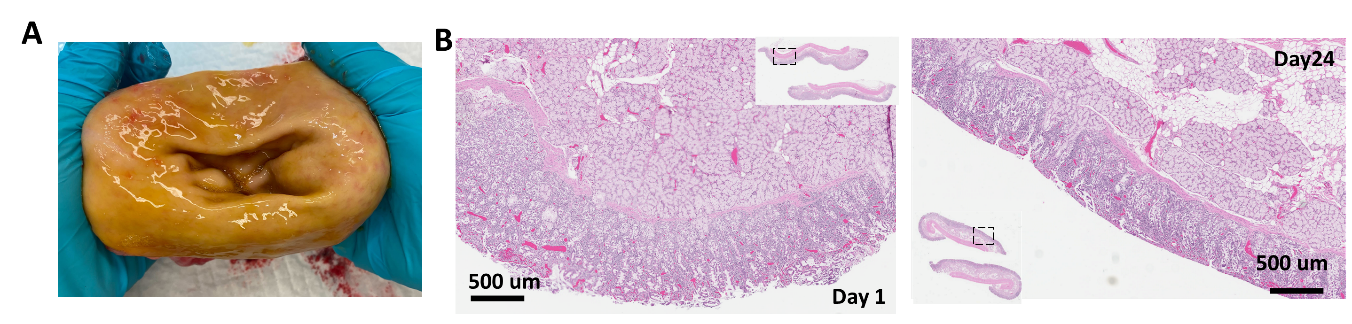

**fig. S16. Long-term mucosa safety 7-60 days after e-GLUE treatment.** **A**) Representative photo of the gastric mucosa 60 days after e-GLUE treatment. **B**) Representative histological results of gastric mucosa 24 days after e-GLUE treatment.

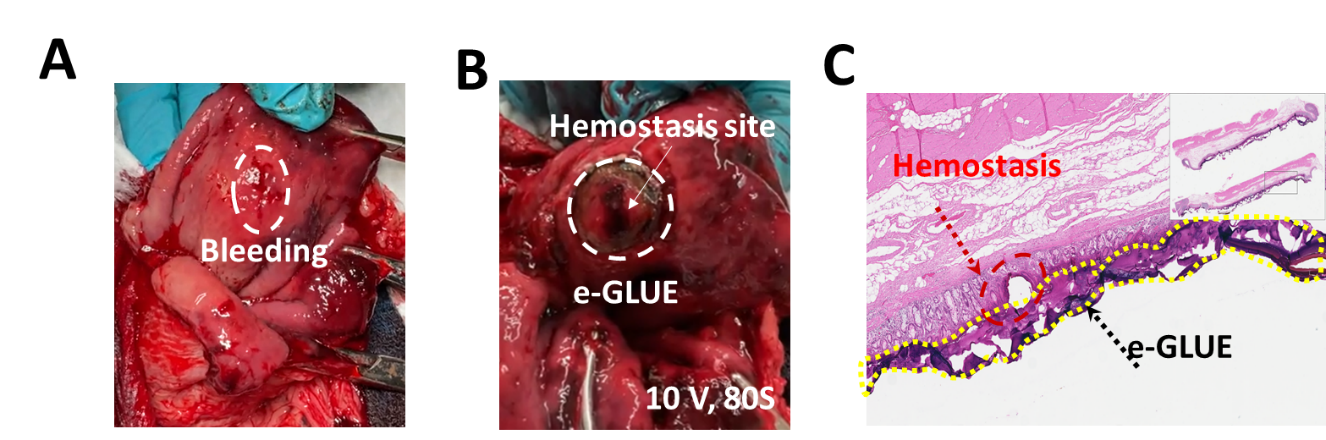

**fig. S17. Representative photos** **A**) before and **B**) after gastric mucosa hemostasis treated by e-GLUE. **C**) The histological result after gastric mucosa hemostasis treated by e-GLUE.

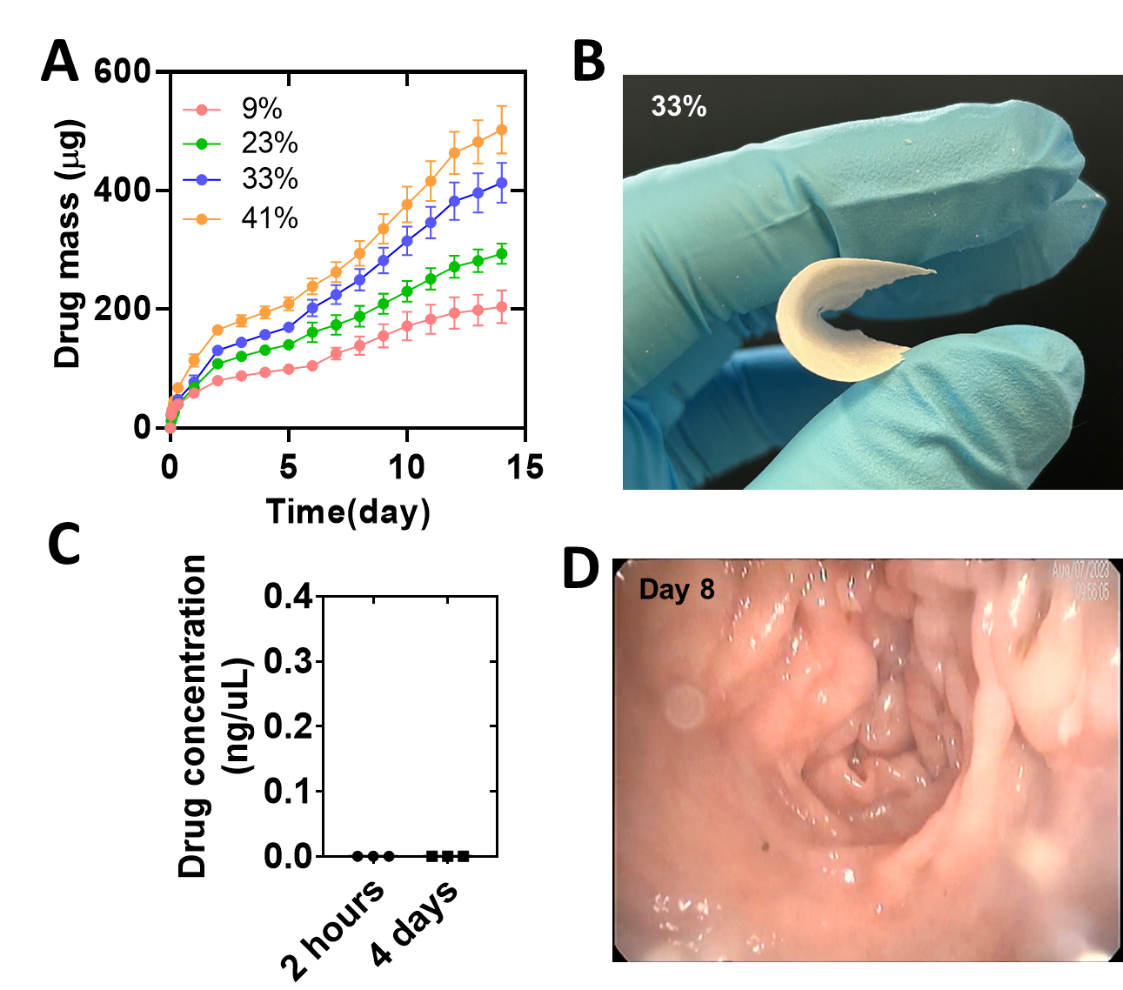

**fig. S18.** **Budesonide-loaded PLGA films as slow-release drug depots**. **A**) Release kinetics of PLGA films with different drug concentrations. **B**) Mechanical flexibility of PLGA films in the drug concentration of 33 wt.%. **C**) Budesonide concentration in the serum over the retention of e-GLUE administration. **D**) The endoscopic view of colon mucosa 8 days after e-GLUE treatment. The values in panels **A** and **C** indicate the mean and the standard deviation (n = 3).

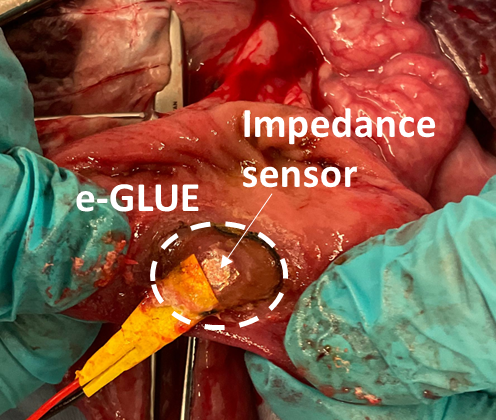

**fig. S19.** e-GLUE enabled a robust interface between intestinal mucosa and the impedance sensors for impedance sensing with enhanced measurement precision.

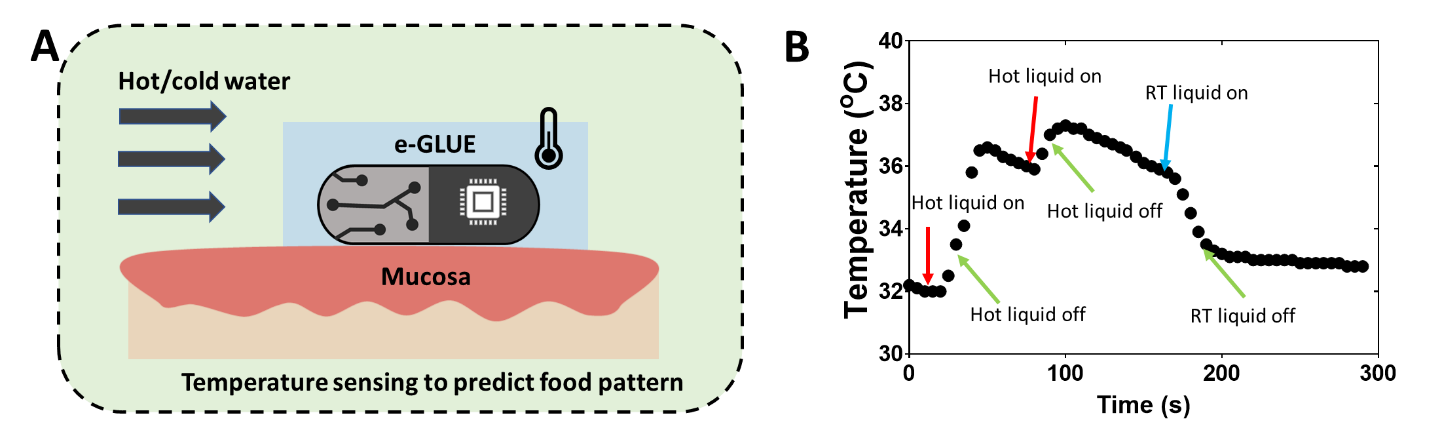

**fig. S20. Localized temperature sensing through stable mucosa-sensor interaction**. **A**) Schematic illustrating the adhesion of an e-GLUE-thermocouple hybrid on mucosal tissues under liquid flow. e-GLUE enabled a robust interface between duodenum mucosa and the temperature sensors. **B**) Localized temperature change during food/liquid administration.
